# Patterns of Microbial Succession and Niche Differentiation Across Depth and Age in a Landfill have Implications for Management Strategies

**DOI:** 10.1101/2025.11.30.686798

**Authors:** Damayanti Rodriguez-Ramos, Liyuan Hou, Patricia Q. Tran, Wanjing Chen, Hans Nielsen-Fox, Fuad J. Shatara, David A. Rivera-Kohr, Joseph H. Skarlupka, Namita P. Shroff, Grace Cagle, Zachary Freedman, Garret Suen, Allison Rathsack, Brian G. Fox, Erica L.-W. Majumder

## Abstract

Despite microorganisms being primarily responsible for landfill material decomposition, limited characterization has been performed across both landfill depth and age. Here, we investigated the microbial communities and physicochemical parameters in two active and two closed landfill wells from surface to bottom, as well as fill dirt and leachate at a sanitary landfill near Madison, Wisconsin, USA. Amplicon community sequencing fungi, bacteria and archaea revealed distinct microbial community structures across landfill sites. The observed patterns of microbial community succession by depth and age mirror the known phases of the landfill life cycle. Younger surface samples were dominated by aerobic fungi, which transitioned to fermentative bacteria and methanogenic archaea in older, deeper, layers. Simultaneously, high species richness was preserved across landfill ages, while reduced evenness at specific depths support spatial niche differentiation. In conjunction with the lack of trends found for physicochemical parameters by depth, this supports niche differentiation driven by the highly heterogenous waste inputs. This study provides the first comprehensive vertical profile of bacterial, fungal, and archaeal communities across landfill depths and ages, highlighting the influence of these parameters on physicochemical factors and microbial distribution within landfills. These results have implications for improving landfill management, including renewable gas energy production, minimizing emissions, and increasing degradation rates.

## 1. INTRODUCTION

Generation of municipal solid waste (MSW), also known as trash or refuse, has grown substantially in recent decades in the United States (U.S.), reaching 292.4 million tons in 2018 or approximately 5 pounds of waste generated per person each day. Of this, 146 million tons (50%) were disposed in landfills (EPA, 2024). With the implementation of government regulations, landfills have evolved into highly engineered and carefully managed facilities designed to safely contain vast amounts of waste. However, decomposition rates are limited by microbial activity that is too slow to keep pace with inputs. This challenge is further compounded by issues with odors and other emissions generated by microorganisms (e.g. methane, ammonia, hydrogen, etc.). Current landfill management practices typically do not consider gas emissions by microorganisms beyond a cursory prediction of methane generation. Therefore, understanding the microbiology of landfills is essential for sustainable waste management.

Landfills are highly complex environments due to their diverse waste inputs, as well as the structure and placement of waste. MSW includes a diverse mix of materials from a variety of sources including households, restaurants, hospitals, and schools. These inputs comprise biodegradable (e.g., food, plants, yard waste) and non-biodegradable (e.g., petro-chemical plastics, metals, glass) items (Godbole et al., 2023). Many landfills also contain other waste types such as biosolids, sewage sludge, construction and demolition debris (C&D), and industrial byproducts. Each material adds different chemicals and elements to landfills. Landfills can extend up to 100 meters below ground and rise as high as 50 meters above the surface (Meyer-Dombard et al., 2020). Over time, landfills can be categorized into "active" and "closed" sites based on their operational status. Active landfills are facilities that are currently receiving waste, whereas closed landfills are those that have reached their maximum capacity and no longer accept waste. Waste is placed in designated “daily cells,” which are specific open sections in the landfill where it is deposited and tightly compacted to reduce space. In addition to waste containment, landfills must manage byproducts of microbial activity, particularly landfill gases and leachate. (Godbole et al., 2023; Meyer-Dombard et al., 2020; Mondal et al., 2023; Sekhohola-Dlamini & Tekere, 2020; Weaver et al., 2019). Once a landfill reaches its maximum capacity, it receives a final cover system, typically vegetative (Mondal et al., 2023). Click or tap here to enter text.

Landfills are dynamic environments where microbial communities play a critical role in the decomposition of waste. Microbial biodegradation processes such as hydrolysis, fermentation (acidogenesis and acetogenesis), and methanogenesis take place in landfills with the output of one process feeding into the next (Sekhohola-Dlamini & Tekere, 2020).

Over time, as waste decomposes due to these microbial and abiotic processes, landfills transition through several distinct biological phases: aerobic (phase 1), anaerobic acid (phase 2), accelerated methanogenesis (phase 3), decelerated methanogenesis (phase 4) and oxygen infiltration – final maturation (phase 5). In full-scale landfills, multiple phases can occur simultaneously due to the heterogeneity of waste and dispersity of cell filling at different times. (Meher et al., 2014; Sekhohola-Dlamini & Tekere, 2020).

Each landfill site is made up of leachate (liquid) and solids (refuse/waste). One of the primary methods for understanding the phases and microbiology of a landfill is through leachate analysis. Leachate is water that has percolated through all the layers of the landfill and reflects an average condition across the site. Research on bacterial communities in leachate has shown that the most predominant phyla include *Gammaproteobacteria, Firmicutes, and Bacteroidetes* (Hirakawa et al., 2025; Song et al., 2015b). Research on archaeal members in leachate has found dominant taxa from the phyla *Euryarchaeota, Halobacterota,* and *Thermoplasmatota* (Grégoire et al., 2023; Krishnamurthi & Chakrabarti, 2013). A few studies have characterized depth profiles for fungi, with one study reporting dominant taxa within the family *Hypocreaceae,* and the genera *Fusarium, and Aspergillus* (Ye et al., 2020).

Leachate contains distinct microbial taxa not typically found in other environments, reflecting the specific conditions within each landfill site. (Collins-Fairclough et al., 2018; Grégoire et al., 2023; Kasonga et al., 2021; Sauk & Hug, 2022; Stamps et al., 2016). As such, only relying on leachate does not reflect the complete microbial community growing on trash at any given time within a landfill, limiting our understanding of how microbial community succession is related to landfill phases.

To characterize landfill microbial communities associated with refuse, researchers have investigated some types of microorganisms at different depths or types of landfills. For example, sampling of the microbial community in cover soils, ranging from 0-150 cm, demonstrated that microbial composition (bacteria, archaea and/or fungi) at these depths is stratified and varies across depths and ages (Sekhohola-Dlamini et al., 2021; Singh et al., 2022; X. Wang et al., 2017; Zabihollahi et al., 2024; Zainun & Simarani, 2018). Multiple studies have also characterized the landfill refuse microbiome across age and depth, particularly in China (Ke et al., 2022; Li, Han, Zeng, et al., 2022a; S. jia Liu et al., 2019; P. Wang et al., 2022; Y.-N. Wang et al., 2021; S. Xu et al., 2017), Japan (Sawamura et al., 2010), Brazil (Morita et al., 2020), India (A. Kumar et al., 2024; R. Kumar et al., 2021; Szulc et al., 2022), and the USA (Schupp et al., 2020), among many other parts of the world. The most predominant bacterial phyla found across landfill sites include *Proteobacteria, Firmicutes*, *Actinobacteria and Bacteroidetes,* with *Firmicutes* being overwhelmingly the most predominant among these taxa, highlighting their importance in waste decomposition. Archaea play a central role in methanogenesis, breaking down waste under anaerobic conditions to produce methane. Studies that examined archaeal communities across landfill depths and ages (Chen et al., 2003; Dong et al., 2015; Krishnamurthi & Chakrabarti, 2013; S. jia Liu et al., 2019; Song et al., 2015a) report *Euryarchaeota* as the predominant phylum, with *Methanosarcinales* and *Methanomicrobiales* as the dominant classes. In contrast, far less is known about fungal contributions to landfill communities, despite their ability to degrade complex organic materials such as lignocellulose, plastics, and other recalcitrant compounds that are resistant to bacterial breakdown (Munir et al., 2024). Only a handful of studies have reported landfill fungal community composition, with one study reporting that the phylum Ascomycota and the genera of *Fusarium* and *Aspergillus* predominated (Ye et al., 2020).

Landfills represent one of the largest engineered microbial ecosystems, with significant implications for greenhouse gas emissions, climate change, and long-term waste management. Understanding the microbiology of landfill is critical for minimizing or predicting methane production, improving waste strategies, and mitigating environmental impacts. Despite advancements in understanding landfill microbiology, a coordinated examination of bacteria, archaea, fungi and physicochemical parameters across various landfill depths and ages has not been reported, particularly for U.S. landfills. To address this knowledge gap, we measured microbial community composition and structure and physicochemical parameters of four full vertical profiles of boreholes, including two from active and two from closed sites within the same landfill. We analyzed the relationship between abiotic and microbial parameters and revealed insights into the microbial ecology of landfills, particularly with respect to succession. These findings will not only enhance our understanding of landfill microbiomes and their role in waste degradation, but have implications for environmental landfill management practices.

## 2. MATERIALS AND METHODS

### 2.1. Site and landfill description

Samples were collected in May 2021, from the Dane County Sanitary Landfill (DCL, Fig. 1G), a MSW facility located in Madison, Wisconsin, United States (43.045355°N, - 89.263094° W). The Madison region experiences a humid continental climate with distinct seasonal variation. Summers are warm, with average air high temperatures around 82°F (27°C) in July Winters are cold, with average lows near 12°F (-11°C). The region also experiences an average annual precipitation of approximately 37 inches. (Madison Climate Data | Wisconsin State Climatology Office, 2025)

**Figure 1:**
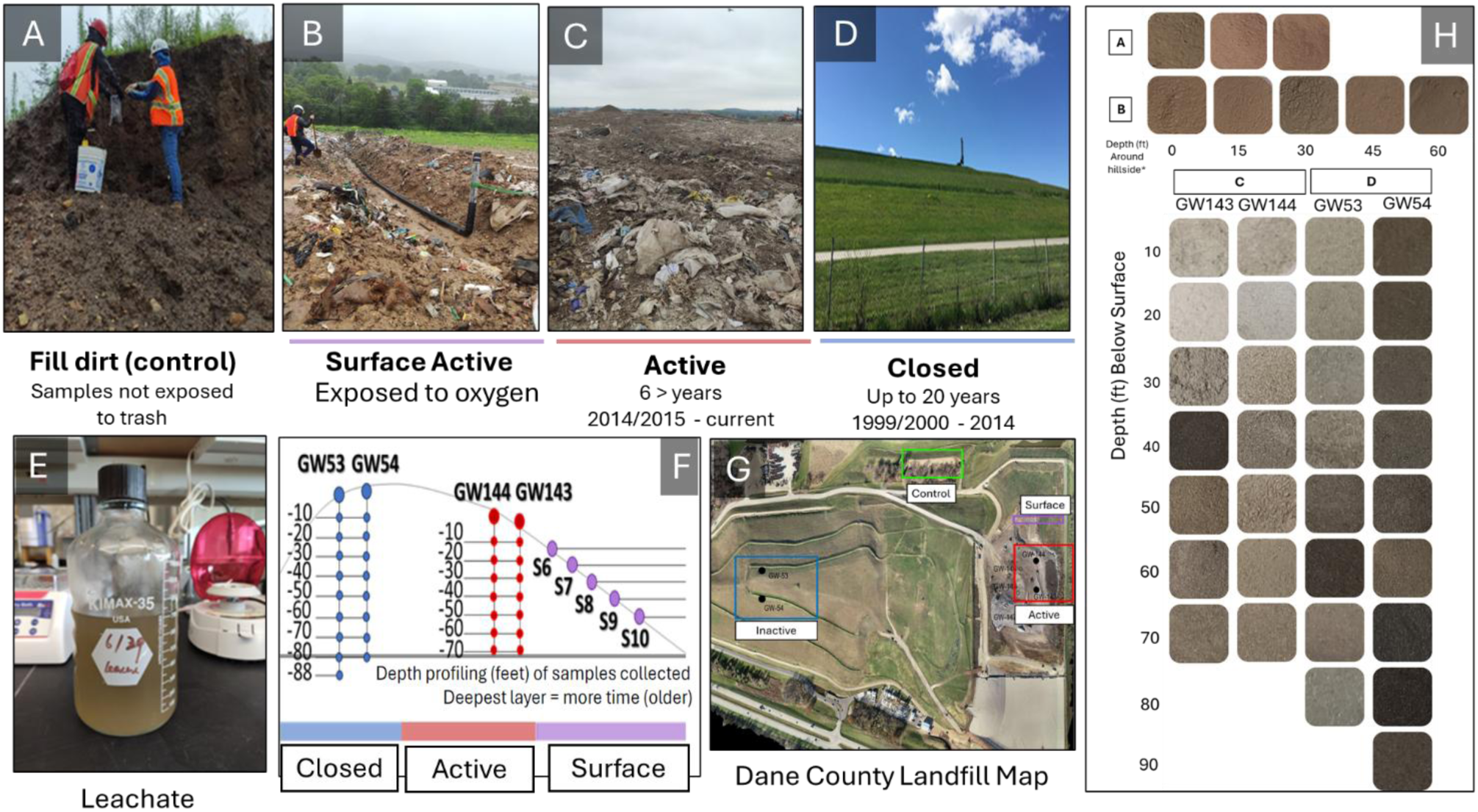
Overview of landfill sampling locations, depth profiling, and soil characteristics. (A) Sampling from fill-dirt (control samples), (B) View of the surface active landfill area, showing exposed waste materials and landfill infrastructure, (C) Active landfill, displaying accumulated waste and operational conditions, (D) Inactive landfill (E) Landfill leachate, (F) Depth profiling diagram showing sampling depths (in feet) at different landfill sites. Deeper layers represent older waste deposits, with sample locations categorized into closed (blue), active (red), and surface (purple) sections, (G) Aerial image of the Dane County Landfill (DCL) site with marked sampling locations, including control (green), surface (purple), active (red), and Closed (blue) landfill sections. The leachate pumpstation is not pictured but is near the main entrance gate, (H) Soil color variations from landfill samples, representing differences in composition, depth, and landfill age.

DCL is a sanitary landfill, meaning that each day’s waste is covered by a few inches of soil and woodchips, and then compacted by machinery into smaller layers in order to preserve space within the landfill. Compacted soil (clay) and plastic liners (polyethylene, PE) are used to protect the surroundings soil and groundwater from untreated MSW, moisture infiltration, and the byproducts of microbial degradation. It has a collection leachate system, and leachate is not recirculated back into the landfill, meaning that this landfill is not a bioreactor landfill that enhances microbial degradation. In addition, the active and closed landfill sites share only one leachate collection system, meaning that the microbial communities of the leachate are a mixture of active and closed sites. (Bioreactor Landfills | US EPA, 2025; Fred Lee et al., 1996) (Fig. S1 and Table S1)

DCL accepts a variety of waste materials, with a focus on ensuring efficient disposal and promoting recycling where possible. Accepted items include MSW, mattresses, box springs, furniture, carpets, stoves, washers, dryers, and some construction and demolition debris that can not be recycled. Tires (without rims) and clean asphalt shingles are accepted specifically for recycling, along with clean branches, logs, and wood chips.

However, to conserve landfill space, easily recyclable or compostable items are restricted. For instance, Freon-containing appliances, propane tanks, cardboard, and aluminum containers are better for recycling. Yard waste, including leaves and grass, is also excluded because they can be composted, with an exception made for invasive weeds. Furthermore, to prevent environmental harm, hazardous materials such as waste oil, toxic chemicals, and electronics are strictly banned from landfill disposal. (Dane County Landfill | Dane County Department of Waste & Renewables, 2025)

To assess microbial and physicochemical changes over time, we used depth as a proxy for relative waste age within each landfill with site status (active vs. closed) representing different landfill aging stages. Given that waste is deposited sequentially from bottom to top, deeper layers are older than surface layers at the same site. At the time of sampling, active sites had been accepting waste for 6 years, while closed sites contained waste deposited over 7 years ago, with the oldest closed site layer being 21 years old. Thus, within each site, depth reflects the relative age of waste, while site status represents the broader chronological context.

### 2.2. Sample Collection

To examine the vertical distribution and impacts of age on the landfill microbial community, we sampled refuse during the installation of the four methane and biogas collection wells in May 2021. We collected samples from two active site wells (GW143 and GW144) and two closed site wells (GW53 and GW54) at depths of every 10 feet (ft) (3 meters) down to 88 ft (∼30 meters) below the surface (Fig. 1F). An auger bucket driller was used to excavate the refuse samples, and a shovel was used to collect samples once the driller dropped the samples. The shovels and buckets were sterilized using bleach between sample collections to avoid contamination. Samples were transferred to sterilized plastic zip-bags, stored on ice in the field, returned to the lab, and frozen at -80°C until further analysis.

We also collected leachate, active surface, and filled dirt (control) samples (Fig. 1E, 1B and 1A). Leachate was collected from all landfill sites to estimate the overall microbial diversity at each site. We obtained approximately 3 liters of leachate directly from the collection tank near the landfill entrance, using sterile glass bottles and buckets. Surface samples (Fig. 1B) were systematically collected from the active hillside of the landfill using bleached shovels and buckets. Sampling began at the top of the landfill (0 ft) and continued down the slope, with samples taken approximately every 15 feet of distance (i.e., at 0 ft, 15 ft, 30 ft, 45 ft, and 60 ft). The sampling area, as shown in Fig. 1B, consisted of a trench around visible landfill infrastructure (e.g., the black drainage tube) with active waste exposure and oxygen availability. Fill dirt (control) samples were collected from piles of soil found within the landfill property, but away from any waste material. This soil was sourced directly from the surrounding area and used by the landfill as daily cover for trash. Since these samples come from local soil that has never been in contact with landfill waste, they serve as a negative control representing the natural background environment.

### 2.3. Physicochemical analysis

Samples were prepared for physicochemical analysis by removing large components of waste material such as stones, glass, plastics, pieces of wood, and other large debris. Samples were dried at 105°C to remove moisture and particles were passed through a 2 mm sieve (No.10), as 2 mm is the US EPA-defined upper threshold of soil particle size (EPA, 1996). After refuse samples were sieved, pictures were taken to record differences in soil color (Fig. 1H).

The pH and electrical conductivity (EC) were determined using a 1:5 (w/v) soil-to-water solution (5 g of soil in 25 mL of Milli-Q water [Milli-Q® Type 1 Ultrapure Water Systems, MilliporeSigma]) (Chukwuma et al., 2021; Sekhohola-Dlamini et al., 2021; X. Wang et al., 2017). The pH was measured using a pH meter (Fisher Scientific), and EC was measured using a Thermo Scientific Orion Star™ electrode. Organic matter (OM) content was determined by Loss on Ignition (LOI) at 400°C for 16 h in a muffle furnace. This process involved measuring 5 g of soils (or refuse) in ceramic cups, placing the cups in the furnace, then measuring the difference of initial weight vs weight after furnace using an analytical balance. The melting of microplastics could have contributed to overall OM, since these melt at high temperatures. Total nitrogen (TN) was determined by using the Total Kjeldahl Nitrogen (organic nitrogen + ammonia-nitrogen) method by USDA/UW-Madison Soils Testing Lab where samples were digested with sulfuric acid and metal catalyst (Nitrogen (Total/Kjeldahl). Metal concentrations, including Ag, As, Cr, Cu, Fe, Mn, Mo, Nb, Ni, Pb, Rb, Sn, Sr, Ti, V, Y, Zn and Zr were quantified using a handheld X-ray fluorescence machine (Innov-X Systems). This measurement was performed on the 2 mm sieved soils. The ions NH₄⁺, NO₂⁻, NO₃⁻, PO₄³⁻, Na⁺, K⁺, Mg²⁺, Ca²⁺, and SO₄²⁻ were measured using the ICS-1100 (Cations) and ICS-2100 (Anions) systems, with quantification performed using Chromeleon software version 7. In this process, 5 g of samples were shaken for 30 min at 120 rpm in Milli-Q water, after this they were filtered using a 0.2 µm filter syringe. Samples were then diluted at 1:100, except active surface samples which were diluted in 1:2 ratio; control samples were not diluted. Dilutions of refuse samples were necessary due to their initially high ion concentrations, which could damage the ion chromatography columns and reduce measurement accuracy, whereas surface and control samples contained lower ion concentrations and therefore required less dilution. Moisture and temperature were measured by DCL employees following their standard procedures.

We were unable to record temperatures directly from GW143 and GW144, so temperature data from nearby wells GW142 and GW145, that were drilled and filled in the same time period, were used. Landfill environmental monitoring data for DCL (database #3018), such as release of oxygen, methane, hydrogen, among others, were obtained from the Waste & Materials Management GEMS on the Web (GOTW) Public Access (https://apps.dnr.wi.gov/gotw/webpages/UserAgreement.aspx). database with wells GW143, GW144, GW53 and GW54 with all parameters and subtypes selected.

### 2.4 DNA extraction, library preparation and sequencing

A total of 39 samples including from active (N=14), closed (N= 17) and surface (N= 5) sites in the landfill, 3 control (fill-dirt) samples (which were homogenized to one sample for analysis), and 3L of leachate were collected (Table S4). Leachate samples were filtered using a 0.2 μm cellulose acetate membrane filters until noticeable build up was formed on the membrane, whereupon they were folded inside a 2 mL PCR tube and stored at -80°C until further extraction. DNA was extracted using the DNAasy Power Soil Kit (QIAGEN), where the first step was modified using 0.4 g of refuse sample or filter, and the Mini BeadBeater Disruptor was used for 2 min, to prevent sample warming and potential DNA denaturation. The concentration and purity of the extracted DNA samples were determined by Qubit fluorometer reagents (Invitrogen, Waltham, MA) and NanoDrop spectrophotometer.

The V4 region of bacterial 16S rRNA, the V6–V8 region of archaeal 16S rRNA, and the ITS2 region of fungal rRNA were amplified via PCR for sequencing on the Illumina MiSeq platform. For bacterial communities, the V4 region of the 16S rRNA gene was amplified using barcoded universal primers with adapters suitable for sequencing on an Illumina MiSeq (V4 Region Primers: F-GTGCCAGCMGCCGCGGTAA, R-GGACTACHVGGGTWTCTAAT; Illumina Adapters: F-AATGATACGGCGACCACCGAGATCTACAC, R-CAAGCAGAAGACGGCATACGAGAT) (Kozich et al., 2013). Archaeal primers targeted the V6–V8 region were used: MiSeq Ar915aF (TCGTCGGCAGCGTCAGATGTGTATAAGAGACAGAGGAATTGGCGGGGGAGCAC) and MiSeq Ar1386R (GTCTCGTGGGCTCGGAGATGTGTATAAGAGACAGGCGGTGTGTGCAAGGAGC), generating ∼610 bp amplicons (Zhou et al., 2017). For fungal sequencing, the ITS2 region was amplified using custom primers (Read 1: TATGGTAATTGTAGCCTCCGCTTATTGATATGCTTAART and Read 2: AGTCAGCCAGCCAACTTTYRRCAAYGGATCWCT), producing amplicons ranging from 450–550 bp (Taylor et al., 2016).

Two rounds of PCR were performed for each target region. In the first round (PCR1), amplification was carried out using region-specific primers. Reactions (25 µL) contained 12.5 µL Kappa DNA polymerase master mix, 0.5 µL of each primer (10 µM), 6.5 µL nuclease-free water, and 5 µL of template DNA (5 ng/µL). Thermal cycling conditions were optimized for each primer set and included denaturation at 95°C for 3 min, followed by 35 cycles of 95°C for 30 s, annealing at a region-specific temperature for 30 s, extension at 72°C for 30 s, and a final elongation step at 72°C for 5 min. Amplification success was verified by gel electrophoresis on a 1% agarose gel with ethidium bromide.

PCR1 products were purified using the Invitrogen PureLink Pro 96 PCR Purification Kit, following the manufacturer’s instructions. Purified products were used as templates for the second round of PCR (PCR2), which incorporated barcodes and Illumina sequencing adapters. PCR2 reactions (25 µL) contained 12.5 µL Kappa DNA polymerase master mix, 0.5 µL (10 nM) barcoding primers (specific to each target), 6.5 µL nuclease-free water, and 5 µL of purified PCR1 product. Thermal cycling included 8 cycles with conditions as in PCR1. PCR2 products were visualized on a low-melt agarose gel, excised, and purified using a Zymo DNA Recovery Kit.

After purification, the amplicons were quantified using the Qubit High Sensitivity Assay and pooled equimolarly to create a final library for sequencing. The pooled library was denatured and diluted to 4 nM using Tris-HCl buffer and NaOH. A 10% PhiX control was spiked into the library to ensure sequencing diversity. Sequencing was performed on the Illumina MiSeq platform using v3 chemistry with 2 × 250bp paired-end reads for archaea and bacteria and 2 × 300 bp paired-end reads for fungi. Contamination prevention measures were employed, including the use of sterile equipment, UV-treated tubes, and negative controls to monitor contamination.

### 2.5 Quantitative real-time PCR (qPCR)

Quantitative PCR (qPCR) was conducted to quantify overall bacterial, fungal, and archaeal communities within samples. DNA was extracted from landfill samples prior to this analysis, and specific primer sets were used to amplify target regions of bacterial, fungal, and archaeal DNA. The primers were validated using reference sequences to ensure specificity for the target organisms.

The qPCR assays targeted specific genetic regions for bacteria, archaea, and fungi, using validated primers (Table S2). We used the 16S rRNA V4 hypervariable region for bacteria, the 16S rRNA V6-V8 region for archaea, and the internal transcribed spacer 1 (ITS1) for fungi.

qPCR reactions were performed in a 96-well plate format using a qPCR machine (CFX Connect Real-Time PCR). DNA samples were diluted to a concentration of 1 ng/µL, and 1 µL of the diluted DNA was used as the template for each reaction.

The qPCR reactions for bacteria and archaea were prepared with a total reaction volume of 12.5 µL, consisting of 0.5 µL of forward primer (F), 0.5 µL of reverse primer (R), 6.25 µL of SYBR Green mix, 4.25 µL of molecular-grade water (DDH_₂_O), and 1 µL of cDNA template. The thermal cycling conditions included an initial denaturation step at 95°C for 5 min, followed by 39 cycles of denaturation at 95°C for 10 s, annealing at 50°C for 30 s, and extension at 65°C for 5 s. A final step at 95°C for 5 s was performed.

For fungal qPCR, the reaction volume was 12.5 µL, including 0.5 µL of forward primer (F), 0.5 µL of reverse primer (R), 6.25 µL of SYBR Green mix, 3.25 µL of DDH_₂_O, and 2 µL of cDNA template. The thermal cycling conditions were : an initial denaturation at 95°C for 5 m, 39 cycles of denaturation at 95°C for 10 s, annealing at 55.6°C for 30 s, and extension at 65°C for 5 s, with a final step at 95°C for 5 s. A no-template control (NTC) and a no-reverse-transcription control (NRT) for each sample were included in the same run. To confirm product specificity, melting curve analysis was performed after each amplification. The number of gene copies per nanogram of DNA (GC/ng) in the sample was calculated using the sample nucleic acid concentration, reported as nanograms per microliter (ng/µL), the volume of sample added to the qPCR reaction (2 µL), and the measured number of gene copies per microliter (GC/µL). Fluorescence data were collected during the extension phase of each cycle. Standard curves for bacteria, archaea, and fungi showed strong linearity, with all assays yielding R^2^ ≥ 0.982. Negative controls, containing all reaction components except template DNA, were included in each run to confirm the absence of contamination.

### 2.6 Sequence Analysis

The resulting bacterial, archaeal, and fungal sequencing files from the Illumina MiSeq were filtered and then analyzed using Mothur v1.47.0. Sequence coverage was estimated during rarefaction analysis and determined using Good’s Coverage, ensuring a minimum of 95% coverage. Archaeal samples ranged from ∼85% to 95% coverage, yet all datasets were filtered prior to downstream analyses. Bacterial and archaeal contigs were aligned against the SILVA database v138 (Quast et al., 2013), while fungal sequences were classified using the UNITE database v8.3 (with a bootstrap confidence cutoff of 80). Pre-clustering was performed using the Needleman–Wunsch alignment algorithm, allowing up to two base pair differences for bacteria and archaea and up to four base pair differences for fungi to minimize sequencing error. Chimeric sequences were detected and removed using the VSEARCH algorithm. Normalization was applied to ensure an equal number of sequences per sample (bacteria: 2180; fungi: 1900; archaea: 214), and samples not shown on graphs were excluded due to insufficient sequencing depth. An operational taxonomic unit (OTU) table was generated for each Domain based on a 97% sequence identity threshold.

Diversity metrics, including Shannon’s diversity, Simpson’s diversity, and Chao’s richness, were calculated. Finally, a distance matrix was generated from each at a 0.03 cutoff to support downstream analyses.

### 2.7 Statistical analysis

Microbial alpha diversity was assessed at the OTU level using normalized abundance data generated in Mothur v1.47.0. Shannon’s (Shannon, 1948), Simpson‘s (Simpson, 1949), and Chao1 (Chao, 1984) indices were calculated, and differences across groups were evaluated with the Kruskal-Wallis (Kruskal & Wallis, 1952) rank sum test and pairwise Wilcoxon tests (Wilcoxon, 1945), with p-values adjusted for multiple comparisons, using RStudio (2024.04.2+764). Non-metric multidimensional scaling (NMDS) was performed using Bray-Curtis dissimilarity and the metaMDS function in the vegan package v2.6-4 (Oksanen et al., 2022) to assess differences in community composition across sampling sites. To emphasize dominant members of the community, low-abundance phyla were collapsed into an ‘Other’ category. Similarly, within each phylum, the most abundant genera were retained, while less abundant taxa were grouped as ‘Other genus’, using ggplot2 and ggnested (Guus Martijn, 2024). Canonical

Correspondence Analysis (CCA) was conducted using the vegan package to explore relationships between environmental variables and microbiome composition. Co-occurrence patterns among taxa were analyzed using an ensemble approach, employing SparCC (Friedman & Alm, 2012) and SPIEC-EASI (Kurtz et al., 2015) as implemented in the SpiecEasi package (Cagle et al., 2024). Top-rank taxa at the phylum and genus levels were visualized using ggnested (Guus Martijn, 2024). Correlations between physicochemical parameters and microbial taxa were visualized with heatmaps generated using the pheatmap package v1.0.12 (Kolde, 2019). Venn diagrams illustrating shared and unique taxa were created using the ggvenn package v0.1.10 (Yan, 2023).

Additional R packages used for data manipulation and visualization included tidyverse (Wickham et al., 2019), readxl v1.4.3, (Neuwirth, 2022), ggtext v0.1.2; (Wilke & Wiernik, 2022), RColorBrewer v1.1-3, (Neuwirth, 2022), cowplot v1.1.1 (Wilke, 2020), plotly (C. Sievert, 2020) colorspace (Zeileis et al., 2020), ggrepel v0.9.3 (Slowikowski, 2023), ggordiplots v0.4.3 (Quensen et al., 2024), and ggpubr v0.6.0 (Kassambara, 2023).

## Data Availability

All sequencing data is available in NCBI BioProject. All other supplementary data are provided in the Supplementary Materials.

## 3. Results

### 3.1 Soil Color Gradients Reflect Depth and Site-Specific Composition in the DCL

The soil samples collected from the DCL showed noticeable variations in color across different sites and depths (Fig. 1H). Surface samples and control (fill-dirt) (Fig. 1B and 1A) exhibited similar reddish and clay tones, which suggests that samples are similar because of minimal waste exposure and/or greater oxygen availability in the surface. Active landfill sites (Fig. 1C) displayed a gradient of lighter to darker tones, reaching the darkest tone around 40 feet in well GW143 before returning to a brown color at lower depths, suggesting multiple degradation states, varied waste composition, locations of transition to anaerobiosis in each well, and differential microbial activity by depth. Closed sites (Fig. 1D) maintained consistent brown and dark tones throughout the sampled depths, suggesting that landfill capping and age play a role in soil stabilization. The leachate sample (Fig. 1E), collected from the landfill pumpstation, represents liquid that has percolated through the waste layers, capturing dissolved organic matter, contaminants, and microbial byproducts. Depth profiling diagrams (Fig. 1F) illustrate the specific sampling depths at each site, highlighting transitions between surface, active, and closed landfill layers. An aerial image of the Dane County Landfill site with marked sampling locations is shown in Fig. 1G, providing spatial context for all sample collection points. Collectively, the panels in Fig. 1 highlight how depth, landfill type, and site conditions drive variability in soil characteristics across the landfill.

### 3.2 Absolute microbial abundance by kingdom varied across landfills sites and depths

qPCR was used to quantify the absolute abundances of microorganisms in each landfill sample by kingdom and revealed variation in microbial community composition and abundance (Fig. 2) that mirrors the pattern described in the five phases of a landfill, which is based on gas production and consumption. Surface samples and the newest and shallowest active site layers tended to have higher microbial abundance (Fig. 2C) and were dominated by bacteria and fungi, with minimal archaeal presence (Fig. 2A&B), reflecting Phase 1 or aerobic processes. Surface samples, such as S6 and S10, also exhibited high microbial overall abundance, with copy numbers reaching 3.34×10^9^ ± 3.25E+08 and 1.53×10^9^ ± 2.68E+08 (Fig. 2C). Active landfill layers, exhibit variable but relatively high microbial copy numbers, potentially likely due to ongoing degradation processes. Overall, the highest total microbial copy numbers were observed in sample 1433 with an overall value of 6.01×10⁹ (Fig. 2C). In active landfill layers (GW143 10-30 ft, GW144 10-20ft), fungal populations were predominant, reaching copy numbers upwards of 5.45×10^9^ ± 5.45E+09 in sample 1431, which was 96% of the total gene copy numbers at that depth (Fig. 2B). With increasing depth and age, fungi decreased while first bacteria, then archaea increased in proportion (Fig. 2A). This suggests that these intermediate layers are in the anaerobic acid phase, and deeper depths are in the rapid methanogenesis phase. Total copy number was generally less in closed landfill layers but some depths (e.g., sample 537 at 3.68×10⁹ ± 7.76E+07) had a higher overall microbial abundance (Fig. 2C). Notably, at 80 to 90 feet, representing approximately 21 years of landfill age, fungal populations showed a slight increase, and archaea had a slight decrease (Fig. 2A). This could be due to landfill age, the oxygen infiltration phase, or these depths are closest to the leachate collection and plastics liner of the landfill which creates a different environment. Leachate samples were also found to also have a high microbial copy number (3.62×10⁹± 1.54E+08), which highlights microbial processes occurring in the liquid systems (Fig. 2C). In contrast to the landfill samples, the control sample (SC) showed a slightly lower microbial abundance, with a total observed copy number of 5.73×10^8^ ± 3.23E+07 (Fig. 2C), which suggests that microbial abundance increases in a landfill relative to the native soil because of the inoculation of microorganisms residing on the refuse materials.

**Figure 2.**
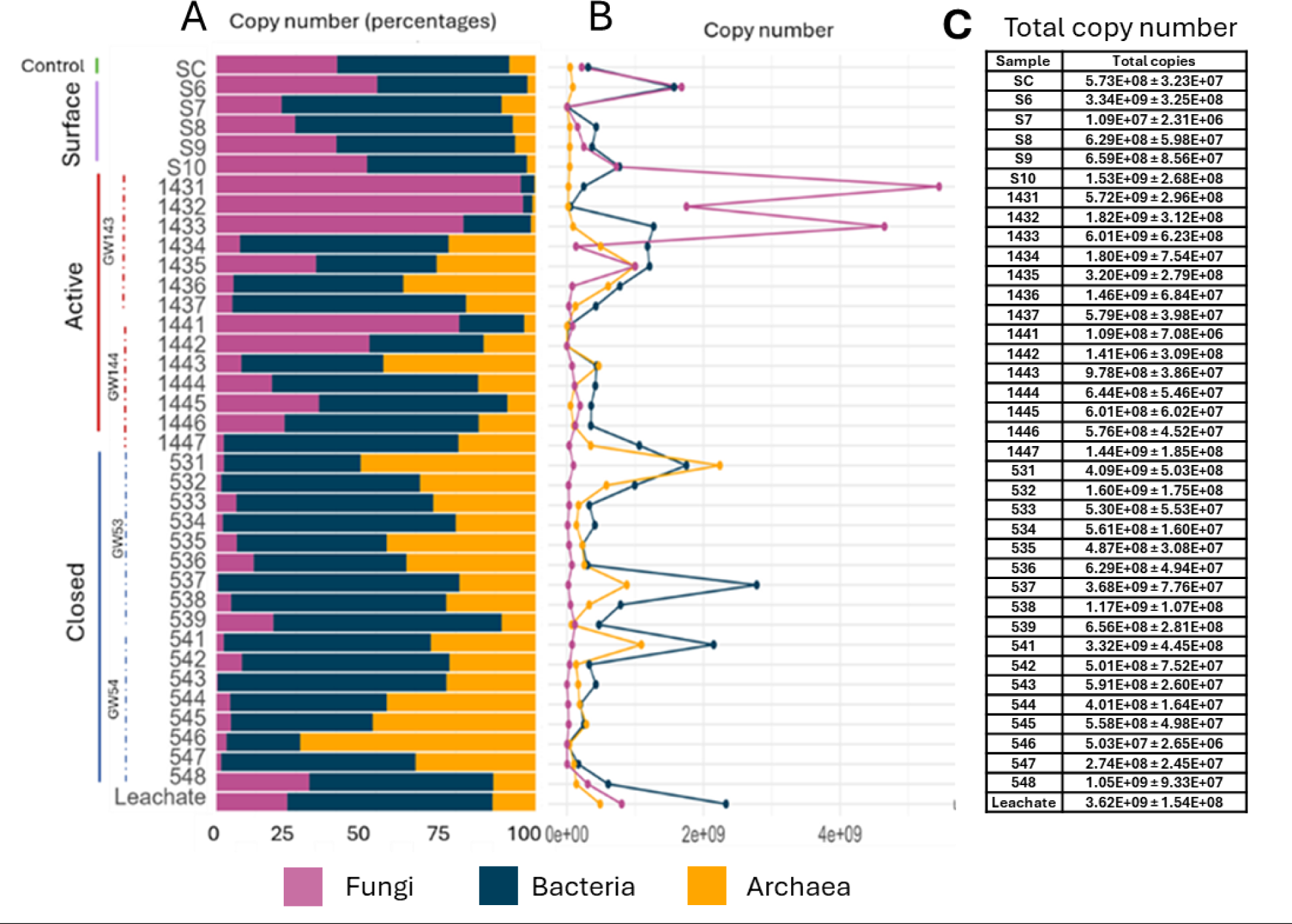
Absolute abundance of microbial kingdoms across landfill sites and depths measured by qPCR. (A) Copy number as percentage (B) Copy number (C) Total absolute abundances per sample

### 3.3 Microbial diversity across landfill sites and depths

Across all domains, a high number of taxonomic groups and OTUs were detected (Table S3), with bacteria showing the highest richness (4,175 OTUs), followed by fungi (2,785 OTUs), and archaea (214 OTUs), reflecting the broad and varied microbial diversity present in landfill environments. The alpha diversity indices by depth reveal notable patterns in microbial diversity across landfill sites (Fig. 3A-C, Fig. S3). Control and surface layers exhibit higher Shannon diversity of bacteria, archaea and fungi, compared to subsurface refuse landfills from active, closed and leachate samples, with average diversity values reaching approximately 5 and 6, respectively (Fig. S3). Active sites showed a high bacterial diversity of more than 4, while closed and leachate samples exhibited lower bacterial diversity, with values around 4 and 2.5 (Fig. 3A). Bacterial diversity was significantly different between active and inactive sites (p-value< 0.05), and between inactive and surface sites (p-value ≤ 0.01) This suggests that subsurface depths have specific microbial niche for the degradation of waste and/or the anaerobic environments, compared to samples that have not been exposed to trash or have more oxygen exposure. Variability or stratification was observed across depths and sites. Each well, either from active or closed sites, and with variable depths did not exhibit a distinguishable increase or decrease in their diversity metrics, For example, samples 1432 (20ft), 534 (40ft), and 537 (70ft), had Shannon’s diversity indices below 3 for bacteria, whereas most of the other samples had values ranging from 3-6 (Fig. 3A). Finally, microbial alpha diversity, as measured using richness estimators (Chao1, Observed OTU’s, ACE) and diversity indices (Shannon, Simpson, Fisher’s Alpha), varied across sample groups for bacteria, archaea, and fungi, suggesting that landfill sites (age), or depth influenced community complexity and evenness (Fig. S4).

**Figure 3.**
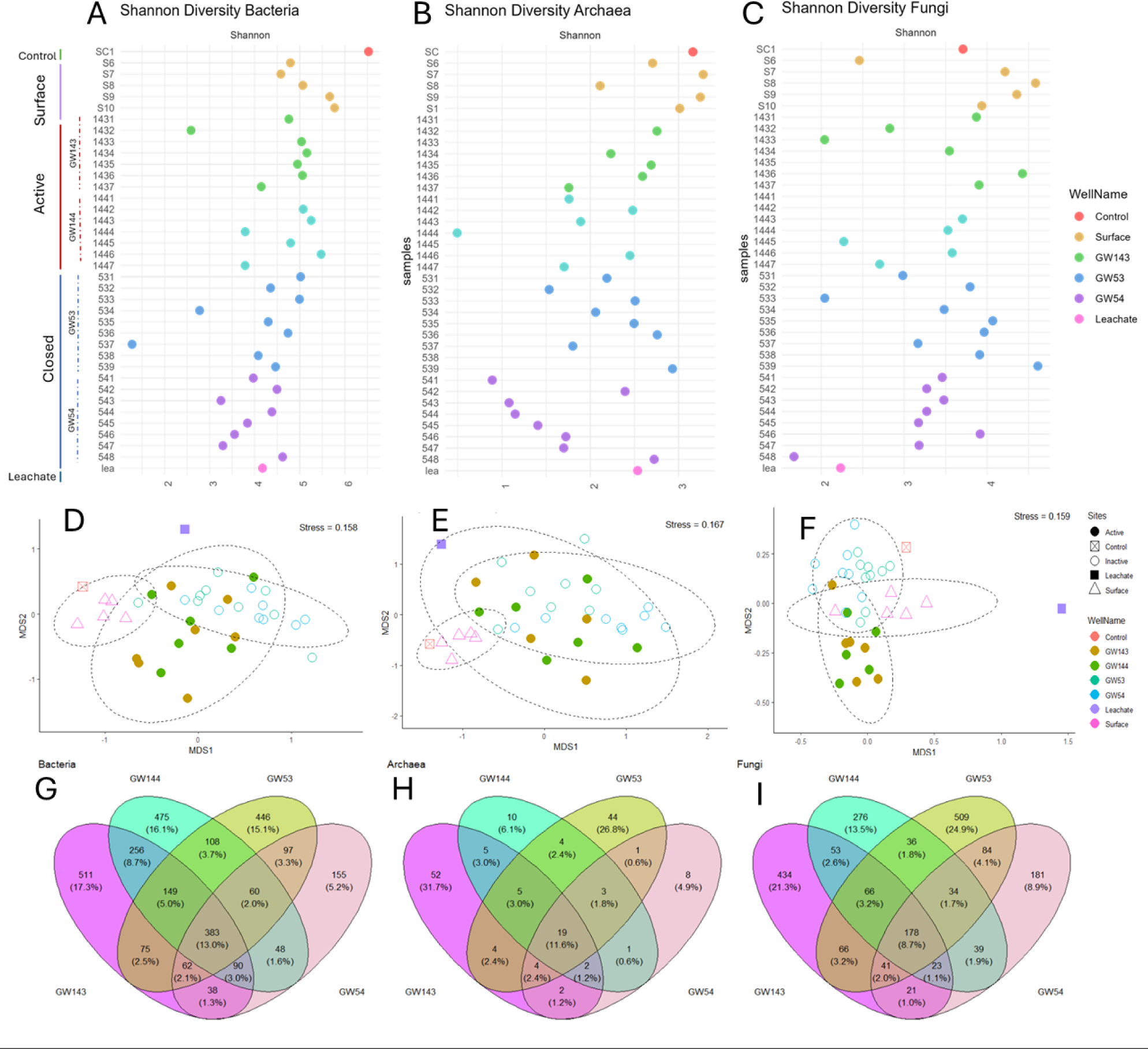
Microbial Diversity Across Landfill Site. Diversity metrics for Bacteria (A, D, G), Archaea (B, E, H), and Fungi (C, F, I). (A-C) Scatterplots depict the Shannon diversity indices from all depths (10–90 feet) and sites. Each point represents the diversity index at a specific depth, with colors indicating the Well or Landfill Site. (D-F) Non-metric multidimensional scaling (NMDS) ordination plots illustrate beta diversity based on Bray-Curtis dissimilarities. Points represent individual samples, colored well and shaped by site category. Calculated dashed ellipses (95% confidence) highlight clustering patterns among site categories. Stress values (<0.2) indicate a good fit of the ordination to the data. (G-I) Venn diagrams display the numbers and percentages shared and unique operational taxonomic units (OTUs) for among landfill wells (GW143, GW144, GW53, and GW54).

Archaeal diversity showed greater variability than bacteria across all depths. On closed sites, well GW54 had Shannon’s values lower than 1.5, while well GW53 ranged from 2-3 (Fig. 3B). This suggests that, in the case of archaea, other factors are involved in diversity rather than refuse age or the sites being closed vs active. Some surface layers exhibited higher archaeal diversity, which could be due to potential ammonia oxidizing archaea (AOA), that can live under oxygenic or low oxygenic environments. This is interesting as archaea are well known for thriving in anaerobic conditions. For archaea (Figure S3B), control and surface sites again showed the highest diversity, followed closely by closed sites. Active and leachate samples had lower archaeal diversity, compared to other sites.

The archaeal communities in between the closed and surface sites were significantly different (p-value < 0.05) These could be due to closed environments (anaerobic), time, or exposure to oxygen. In addition, this suggests that, while archaeal communities in landfill environments are diverse, conditions in less disturbed sites (closed sites) might allow for a broader range of archaea, including methanogens and other specialized groups.

Fungal diversity was also variable across depths, and was higher in surface samples, with most values above 3, and decreases in some depths, as seen in sample 548 (80 ft), where fungal diversity falls below 2 (Fig. 3C). Similarly to archaea, there were different fungal diversities, especially in closed sites of the landfills, with GW53 having very stratified fungal diversities ranging from 1-4+, whereas GW54 exhibited values around 3-3.5. Fungal diversity (Fig. S3C) follows a similar pattern to the bacteria, with the highest diversity observed in surface and control sites, and the lowest in leachate samples. This suggests that environments with higher oxygen availability may support a richer fungal community, potentially involved in the early stages of waste degradation, such as the breakdown of food waste. In contrast, leachate samples exhibit lower fungal diversity (Fig. 3C and S3), indicating that the leachate environment may select for specialized fungi capable of tolerating extreme conditions and metabolizing specific compounds.

By calculating Shannon diversity index by wells instead of by sites, we found variation in microbial diversity across different wells from within the same site. As shown in Figure S3D, the bacterial communities in closed wells GW53 and GW54 were significantly different than the control site with p < 0.05, and p ≤ 0.01, respectively. Archaeal alpha diversity (Fig. S3E) also shows distinct patterns across wells with Shannon’s diversity varying between the same closed well of GW53 and GW54, with GW53 having a lower diversity index (p < 0.05). Similar findings were observed for active wells, with GW143 having a higher diversity, relative to GW144. Statistical differences (p < 0.05) were also observed between GW143 and the leachate; between GW53 and the leachate; and between GW53 and the control.

However, significant differences in fungal diversity (Fig. S3G) were not observed between wells, although slight shifts in diversity within wells were found. Overall, this trend suggests that microbial community diversity (bacteria, archaea, fungi) is influenced by the local site conditions of each well, which can very substantially from well to well within a relatively small landfill area.

Beta diversity, as determined by Bray-Curtis dissimilarity analysis and visualized with NMDS plots, demonstrated distinct clustering patterns for the microbial communities across site categories (Fig. 3D-F). Leachate and control sites harbored unique microbial compositions, while surface and control samples showed overlapping bacterial and archaeal communities, indicating shared taxa. Active and closed sites exhibited partial overlaps for bacteria and archaea, whereas fungal communities were more distinct, reflecting site-specific influences on microbial diversity and composition.

Venn diagram analyses of species richness further highlights the distinct microbial communities within each well (Fig. 3G-I). For bacteria, GW143 and GW144 contain the highest numbers of unique taxa (511 and 475, respectively), while 383 taxa (13.0%) were shared across all four wells. Archaea show minimal overlap, with only 3 shared taxa (1.8%) across wells and GW143 having the most unique taxa (52, 31.7%). Fungi also exhibit higher differentiation, with GW53 containing the largest number of unique taxa (509, 24.9%), and only 178 taxa (8.7%) shared across all wells. This indicated that each site harbors specific taxa, potentially unique to waste input. .

### 3.4 Relative abundance of microbial taxa across the landfill site

#### 3.4.1 Bacterial Community Composition

Bacterial communities exhibited clear differentiation by depth and site type (Fig. 4A). Surface and control samples shared similar taxa, predominantly *Proteobacteria, Actinobacteria, and Acidobacteriota*. Among the control samples, low-abundance genera were present, with the most dominant *being Bacteria_unclassified (Otu00278, 1.29%), KD4-96_ge (Otu00116, 1.53%), and Intrasporangium (Otu00014, 2.10%)*. These patterns could be attributed to the high Shannon diversity and varied functional roles of bacteria in landfill sites.

**Figure 4.**
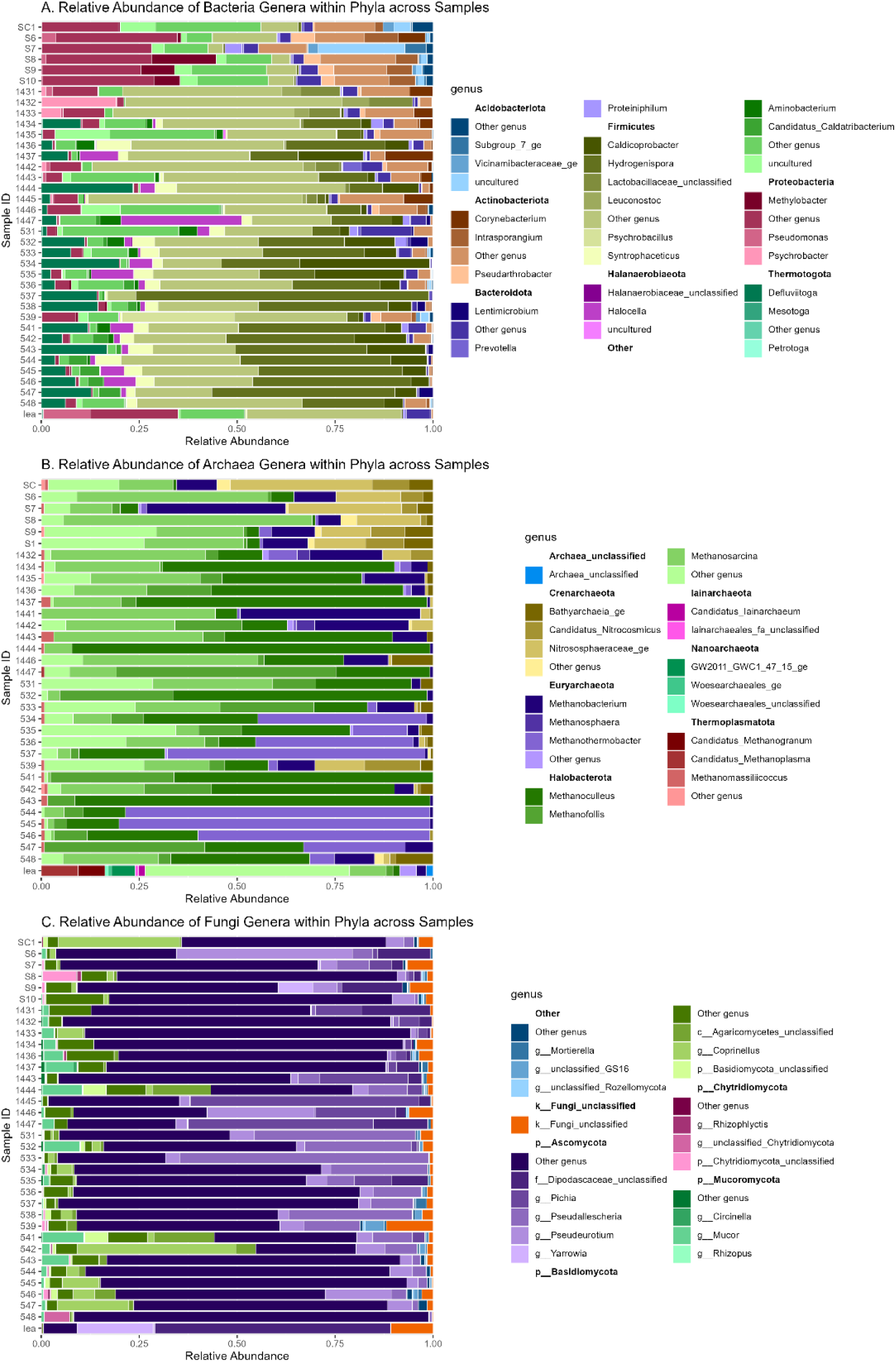
Relative abundance of microbial genera across landfill depths and sites. Stacked taxa bar plots depict the relative abundance of bacterial (A), archaeal (B), and fungal (C) genera grouped by their respective phyla and organized by site type and descending depth.

At surface sites, distinct genera such as P*seudarthrobacter (Otu00019), Acinetobacter (Otu00083), Ellin6067 (Otu00070), uncultured_ge (Otu00039, 16.06%), Intrasporangium (Otu00014), and Methylobacter (Otu00043)* were identified. *Pseudarthrobacter*, previously detected in landfills, plays a crucial role in plastic degradation processes (Chung et al., 2022). *Intrasporangium*, a Gram-positive aerobic bacterium, has also been noted for its potential nitrate reduction under anaerobic conditions (Amachi et al., 2024). Notably, the presence of aerobic methane-oxidizing bacteria such as *Methylobacter* is significant, since they utilize methane as an energy source, potentially reducing landfill methane emissions (Tveit et al., 2023). Their occurrence at surface levels is expected, given their role in aerobic degradation.

In the active surface sites, *Firmicutes* were present in low abundance but increased significantly with depth, in active and closed landfill sites, suggesting a key role in waste degradation in anaerobic environments. Genera such as *Lactobacillaceae_unclassified (Otu00017)*, *Levilactobacillus (Otu00048)*, *Psychrobacter (Otu00056)* and *Carnobacterium Otu00028)* were the most abundant at depth from 10-30 ft from active sites wells GW143 and GW144, suggesting key fermentation processes at these depths, and landfill phases such as anaerobic acid. At 40-70 ft, the phylum *Thermotogota* was present, with certain depths being predominated by the genus of *Defluviitoga*. The presence and predominance of these taxa further indicate the development of anaerobic conditions at these depths, possibly reflecting a shift toward later stages of anaerobic degradation in the landfill.

In active sites such as GW143 (10–30 ft depth), dominant genera included *Levilactobacillus (Otu00048), Lactobacillaceae_unclassified (Otu00017), Corynebacterium (Otu00012), and Psychrobacter (Otu00056)*. *Lactobacillaceae* taxa, particularly *Levilactobacillus and Corynebacterium*, are fermenters, producing lactic acid, acetic acid, and other metabolites. These taxa have been reported in various landfill studies (Chukwuma et al., 2021; S. Xu et al., 2017). Research on refuse decomposition in a French landfill found that *Cellulomonas, Microbacterium,* and *Lactobacillus* predominated in fresh, 1-year-old, and 5-year-old refuse samples (Pourcher et al., 2001). Although *Lactobacillus* can be associated with the aerobic phase of landfill decomposition (S. Yang et al., 2021), other studies have shown that certain species can ferment glucose and xylose into lactic acid under anaerobic conditions and low pH (Chukwuma et al., 2021; Cubas-Cano et al., 2019) aligning with the acidic pH of samples at these depths.

*Psychrobacter* (Otu00056) was also identified in our landfills, and previous depth-profiling studies have detected this genus in upper or middle landfill layers (Li et al., 2022; Y.-N. Wang et al., 2021). Its presence may be linked to refuse age, as studies from a Chinese landfill reported *Psychrobacter* in 0–1-year-old waste (Y.-N. Wang et al., 2021). However, as landfill depth increases, oxygen depletion and rising temperatures limit the growth of *Psychrobacter*, despite their strong ability to degrade organic compounds, (Li et al., 2022) which could be a reason they decrease in our sites. Furthermore*, Psychrobacter* species have been implicated in microplastic degradation (Ou et al., 2024).

In GW144 (10–30 ft depth), similar taxa were observed as in GW143, with the addition of *Turicibacter (Otu00086), Romboutsia (Otu00108), SBR1031_ge (Otu00094), and Spirochaetaceae_unclassified* (Otu00051). *Turicibacter*, another member of the gut microbiota, is an anaerobic, spore-forming genus likely involved in the heterotrophic degradation of organic matter through polysaccharide fermentation (Vilajeliu-Pons et al., 2016), while *Romboutsia* is an anaerobic fermenter. The predominance of these taxa suggests an active anaerobic acid phase, characterized by a pH of 5.5–6.5, aligning with the acidic conditions observed in both active wells. *SBR1031* has been identified in anaerobic digestion processes, particularly in the degradation of wood vinegar wastewater (Hua et al., 2020). Members of this group encode key genes for acetogenic dehydrogenation, enabling the utilization of ethanol as a carbon source (Xia et al., 2016). *Spirochaetaceae*, which includes both aerobic and anaerobic species, has been reported in landfills and plays a role in lignocellulose degradation (Ransom-Jones et al., 2017).

At greater depths (40–70 ft), Firmicutes remain prevalent, with additional phyla appearing, including *Thermotogota, Spirochaetota, Chloroflexi, Synergistota, and Halanaerobiaeota. Thermotogota* is characteristic of landfill environments and has been previously reported at these depths. Notably, temperatures at these depths can reach up to 140°F, which supports the growth of thermophilic taxa. Key genera identified at these depths also include *Corynebacterium (Otu00012, Otu00106), Desulfosporosinus (Otu00047), Cryobacterium (Otu00046), Defluviitoga (Otu00001), Hydrogenispora (Otu00004), Aminobacterium (Otu00010), Caldicoprobacter (Otu00002), and Halocella (Otu00013)*.

At 40 ft in GW144, *Defluviitoga (Otu00001)* comprised 23.38% of the community. This genus has been previously found in deep 40m landfill layers (Ke et al., 2022). Its growth may be stimulated by increasing temperatures, as it thrives in thermophilic anaerobic conditions, effectively degrading organic materials, metabolizing sugars, and producing hydrogen (Ke et al., 2022). Other dominant taxa included *Haloimpatiens* (Otu00065, 11.33%), which has been isolated from contaminated environments such as wastewater and plays a role in formate, acetate, lactate, and ethanol fermentation and lignin degradation (Wu et al., 2016; Zhu et al., 2022). Additionally, *Caldicoprobacter* (Otu00002, 8.52%) and *Bifidobacterium* (Otu00069, 5.29%) were present, with the latter known for its carbohydrate metabolism and adaptations to landfill conditions (Ventura et al., 2016).

Previous studies have linked *Bifidobacterium* abundance in landfills to increasing depth, alongside genera such as *Rhabdanaerobium* and *Saccharopolyspora* (Shao et al., 2023).Click or tap here to enter text.Additionally, *Weissella* (Otu00042, 3.86%) belongs to the lactic acid bacteria family and plays a key role in food fermentation, producing lactic acid and polysaccharides. This genus also exhibits resistance to various antibiotics (Wan et al., 2023). In high-temperature landfill environments, *Weissella* and *Lactobacillus* are more abundant, as they are facultative anaerobic bacteria capable of fermenting glucose and other sugars to produce lactic acid (Shao et al., 2023). At 60 ft, key taxa included *Hydrogenispora* (Otu00015) and *SBR1031_ge* (Otu00060). *Hydrogenispora* is an obligate anaerobe capable of hydrogen production and glucose fermentation, yielding acetate, ethanol, and hydrogen as end products (Y. Liu et al., 2014). At 70 ft, the dominant genera *were Halocella (Otu00024), Hydrogenispora (Otu00006), and Treponema (Otu00029)*.

*Halocella*, a halophilic bacterium, has been extensively reported in landfill studies (Song et al., 2015b). The presence of *Treponema*, an anaerobic spirochete, is notable, as it has been found in landfill depths of approximately 2 m (Li et al., 2022).

In closed landfill sites, bacterial communities were distinct and dominated by *Firmicutes,* particularly the genera *Hydrogenispora.* Other taxa such *Defluviitoga,* and *Caldicoprobacter* were also dominant at these sites. This suggests key taxa involvement in methanogenesis and/or fermentation under anaerobic environments. Leachate samples contained dominant taxa such as *Pseudomonas (Otu00073), Exiguobacterium (Otu00058),* and *Planococcaceae_unclassified (Otu00035)*. In addition to the dominant genera, a substantial portion of the bacterial community was composed of low-abundance taxa (<10% relative abundance), as shown in Figure S2A. Although individually rare or low-abundant, these taxa collectively contribute to the overall diversity and functional potential of the landfill microbiome, highlighting the presence of many genera that persist at low levels across different landfill environments.

#### 3.4.2 Archaeal Community Composition

The archaeal community structure also varied significantly across landfill depths and site types (Fig. 4B). In surface and control samples, the predominant phyla were *Crenarchaeota* and *Halobacterota*, with abundant genera such as *Nitrososphaeraceae* and *Candidatus Nitrosocosmicus*, suggesting common soil archaeal taxa and nitrogen cycling processes. In active sites, the archaeal community shifted, with a decrease in *Crenarchaeota* and an increase in *Halobacterota*. Prominent methanogens included *Methanoculleus* (e.g., Otu00002 in GW143 at depths 1432-20 ft and 1437-70 ft) and *Methanosarcina* (Otu00008). In closed wells, the archaeal community was dominated by thermophilic and methanogenic genera, such as *Methanothermobacte*r (Otu00003), which clearly dominated at depths between 40 and 70 ft. This pattern shows that closed sites, due to long-term anaerobic conditions resulting from capping and aging of the refuse, harbor distinct archaeal taxa adapted to these environments. Leachate samples exhibited a unique archaeal composition, including phyla *Iainarchaeota, Thermoplasmatota,* and *Nanoarchaeota*, with dominant genera *Methanocorpusculum (Otu00012), Methanosaeta (Otu00011),* and *Candidatus Nitrosocosmicus (Otu00016).* Similarly, as with bacteria, Figure S2B shows that archaeal communities also include a high proportion of low-abundance taxa (<10% relative abundance), particularly in surface and leachate samples, suggesting functional specialization in response to these conditions. This distinct composition likely reflects the highly specialized and variable microenvironments within leachate, supporting a diverse range of archaeal lineages, including those involved in methane production and nitrogen cycling.

#### 3.4.3 Fungal Community Composition

The fungal community composition (Fig. 4C) across all landfill sites was dominated by *Ascomycota*. In active sites, layers from 10-30 ft, well GW143 exhibited a high relative abundance of *Penicillium (Otu00006)* and *Teunomyces sp. (Otu00012)*. In GW144, *Pichia fermentans* was prominent, suggesting key fermenters at these initial layers, and contributing to breakdown of components for other microorganisms. In closed wells, the fungal community shifted toward anaerobic and stress-tolerant fungal taxa. Well GW53 was dominated by *Pseudallescheria boydii (Otu00004)* and members of the family *Didymellaceae*, while well GW54 featured *Alternaria destruens* and *Tricellula* (Otu00009). One curious fact is that both wells GW53 and GW54 came from same age or closed site, however their relative abundance of fungal taxa such as *Pseudallescheria boydii* was different, contrary to what we have seen in bacteria and archaea, wherein closed sites harbored the same/similar taxa. This suggests that fungal taxa may depend on a specific waste or unique environment. In sample 548, *Cladosporium (Otu00016)* accounted for approximately 60% of the fungal community. Leachate samples displayed a distinct fungal composition, with unclassified members of the family *Dipodascaceae (Otu00049)* comprising around 60% of the community. Another significant genus in leachate was *Yarrowia lipolytica*, which represented about 15% of the fungal community, suggesting taxa with diverse metabolic capabilities can survive in these environments. As with bacteria and archaea, fungal communities exhibit a large proportion of low-abundance genera (Figure S2C), highlighting the presence of rare but potentially important taxa contributing to waste degradation or stress tolerance.

### 3.5 Concentrations of physicochemical parameters of refuse by depth and sites

Physicochemical measurements of landfill samples were highly variable across landfill sites and did not have a clear trend linked to depth (Figure 5 and Table S4). Control and surface-active samples had an alkaline pH with average of 8 (Fig. 5), low EC of <1 (Table S4), and relatively lower concentrations of all other measured factors compared to subsurface active and closed sites of landfills. Active sites showed variation of pH levels where certain depths had an acidic pH of 5 whereas other depths had neutral pH of 7 (Fig. 5 pH, green & cyan lines). Active sites had higher EC (mS/cm) values up to 6.69, closed sites had a value up to 4.58, surface-active 0.05 and control 0.09.

**Figure 5.**
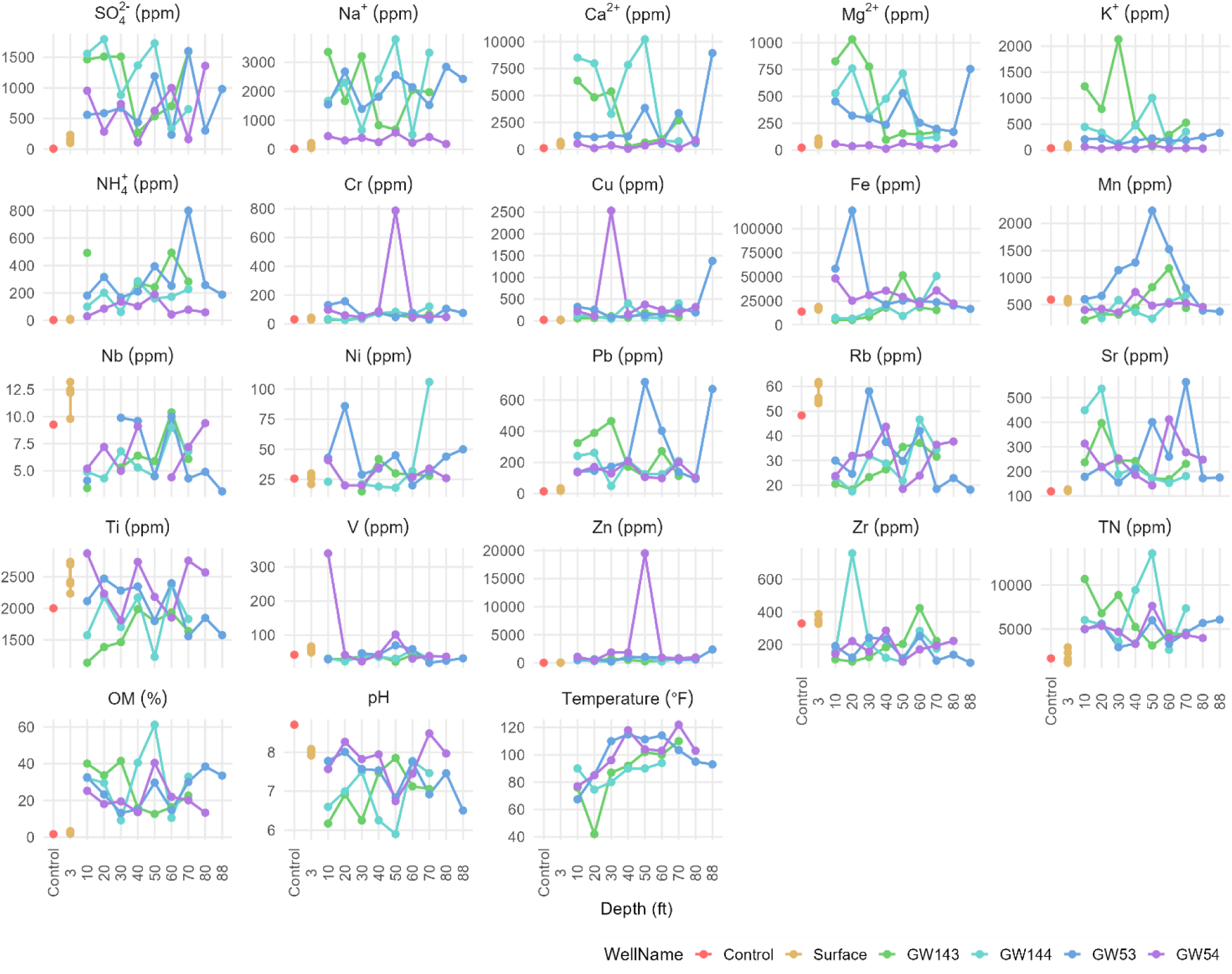
Physicochemical variations across landfill depths. Line graphs represent the trends of physicochemical parameters including cations (ppm), anions (ppm), heavy metals (ppm), temperature (°F), pH, total nitrogen (TN - ppm), organic matter (OM - measured as % weight loss by the ‘loss on ignition’ method). Different colored lines represent distinct landfill sites, as indicated in the legend. The *X*-axis represents sample depth, while the *Y*-axis represents concentration levels in respective units.

Organic matter and total nitrogen levels were relatively high throughout both active and closed sites of the landfill, though substantial variability was observed (Table S4)- Active wells ranging ∼9%–61% in OM, and TN ∼2643 to 13,584 ppm, closed wells ranging ∼13.2% to 40.5% in OM, and ∼2932 to 7631 ppm in TN.

Comparatively, closed sites still showed high concentrations of the physicochemical parameters, with the exceptions of slightly lower organic and ion content, possibly due to stabilized conditions over time. The observed trends indicate sufficient nutrients to support a range of microbial metabolism but with spatial variability in elemental distribution and landfill chemistry across depths.

Within the installations of methane gas wells, collection of data such as methane, oxygen and temperature was recorded. Data has been collected from 5/24/2021 up to 9/13/2024 (Table S5). Gas composition and temperature measurements collected from four gas wells at DCL showed clear patterns indicative of active anaerobic decomposition processes (methane production). Across all wells and sampling dates (meaning of gas measured), methane concentrations remained high, ranging between 47.8% and 53.4% (vol %). Both active and closed wells exhibited similarly elevated methane levels, suggesting that methane is being produced in both active and closed sites.

Oxygen concentrations were consistently at 0.0% (vol %) in most of the wells and time points, however there were some exceptions that oxygen ranged from 1.4-3.7% in wells GW53 and GW54 (Table S5). These oxygen fluctuations were not associated with changes in methane production or temperature. This absence of oxygen highlights the anaerobic environments, which supports methanogenic activity across both active and closed sites. Temperature readings varied from 105.8°F to 114.0°F (40°C to 46°C), remaining stable across all wells. These elevated temperatures fall within the optimal range for mesophilic and thermophilic microbial populations, indicating ongoing microbial degradation and methane processes. While no significant differences in methane, oxygen, or temperature values were observed between active and closed wells, the continued presence of anaerobic conditions and high methane levels in the closed wells suggests that microbial activity and methane generation persist long after landfill closure.

To explore how these physicochemical gradients influence microbial community composition, we performed Canonical Correspondence Analysis (CCA) using the top 20 genera from bacterial, archaeal, and fungal datasets (Figure S5). The CCA plots revealed distinct clustering patterns, indicating that microbial taxa are differentially structured by environmental variables such as pH, ammonia, and sulfate. This highlights the influence of landfill chemistry on shaping domain-specific microbial communities and supports the presence of fine-scale niche partitioning driven by chemical gradients.

### 3.6 Certain physicochemical factors are correlated with taxa

The heatmaps illustrate the relationships between microbial genera and physicochemical parameters across active and inactive landfill sites (Fig. 6). The bacterial community exhibited distinct correlations with environmental parameters in both active and inactive landfill sites. In active landfill sites, bacteria showed diverse associations with physicochemical conditions (Fig. 6A). For example, EC was significantly positively correlated with *Unclassified Spirochaetaceae, Methylobacter, Unclassified Anaerolineaem, Methylocystis, Unclassified Hungateiclostridiaceae, Crenothrix, Vicinamibacteraceae_ge, Intrasporangium* and *KD4-96_ge. Aminobacterium* displayed a strong positive correlation with Mn and *Lentimicrobium, Alkalifexus* and *Candidatus Caldatribacterium* exhibited a positive association with Ni, which may indicate metal tolerance or utilization; *Unclassified Lactobacillaceaea* showed strong positive correlations with Mg^2+^. In contrast, bacterial communities in inactive landfill sites displayed more stable correlations with environmental parameters (Fig. 6B). For example, Ca²⁺ was positively associated with multiple genera, including *Limnohabitans, KD4-96_ge, Cryobacterium*, and *Pseudarthrobacter*, indicating a preference for calcium-enriched environments or that Ca is needed as an enzymatic co-factor. *Ruminiclostridium* exhibited a strong positive correlation with NH₄⁺, suggesting a role in nitrogen transformations. Additionally, *Weissella, Leuconoc* and *Unclassified Lactobacillaceae* were significantly positively associated with V and NO_3_^-^.

**Figure 6.**
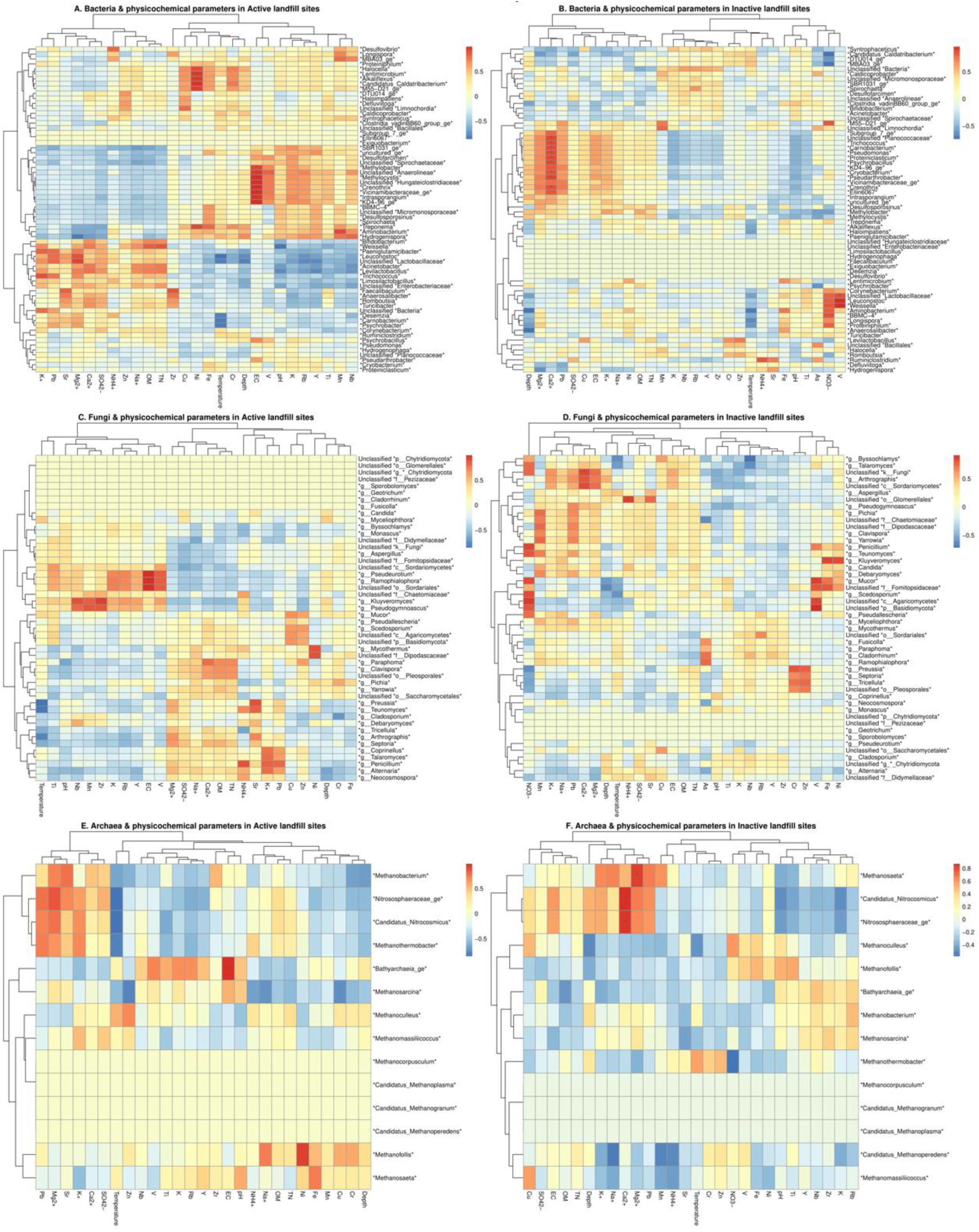
Correlation of microbial genera with physicochemical parameters in active and inactive landfill sites by kingdom. Panel (A-B) bacterial, (C-D) archaeal, and (E-F) fungi. Columns represent physicochemical parameters, while rows represent microbial genera, clustered based on their relative abundance and response to environmental variables. The color scale indicates the strength of the Pearson correlation: red for strong positive correlations, blue for strong negative correlations, and yellow for no significant relationships.

Archaea also displayed distinct environmental associations depending on landfill operational status (age). In active landfill sites, archaeal communities exhibited varied correlations with physicochemical parameters, reflecting the dynamic conditions of waste degradation (Fig. 6C). *Methanobacterium, Methanothermobacter, and Methanosaeta* were positively correlated with pH, EC, and NH_₄_⁺. Strong correlations with *Methanosaeta* and *Methanosarcina* with organic matter (OM) and total nitrogen (TN) were also observed. In case of metals, *Methanofollis* and *Candidatus Methanoperedens* show a notable correlation with Fe and Mn. Nitrogen Cycling Archaea such as *Nitrososphaeraceae_ge* and *Candidatus Nitrosocosmicus* (ammonia-oxidizing archaea) correlate with NH₄⁺ and TN, indicating their role in nitrogen cycling, likely through ammonia oxidation. *Bathyarchaeai* has a strong significant positive correlation with EC. In closed landfill sites, archaeal communities demonstrated more stable associations with specific environmental parameters (Fig. 6D). *Methanosaeta* showed positive correlations with Ca²⁺, Mg²⁺, and NH₄⁺. *Candidatus Nitrosocosmicus* and *Nitrosospharaeraceaea* exhibited strong positive associations with Ca²⁺. *Methanothermobacter* had a positive correlation with temperature, demonstrating its thermophilic nature.

In active landfill sites, fungal genera such as *Pseudeurotium, Ramophialophora, Unclassified Sordariales* were positively correlated with EC (Fig. 6E). Others were positively correlated with metals (Nb, Mn, Zr) such as *Kluyveromyces and Pseudogymnoascus*. Other taxa such as *Coprinellus, Talaromyces, Penicillium, Alternaria and Neocosmospora* were correlated with K and Pb. In inactive landfill sites, several fungal genera, including *Mucor, Unclassified c_Agaricomycetes* and *Unclassified p_Basidiomycota* had significant positive correlations with V (Fig. 6F). Unclassified o_Glomerales was significantly positively correlated with NH4^+^. Multiple taxa were positively correlated with NO3^-^ such as *Byssoclamys, Talaromyces, Unclassified Fungi, Penicillium, Teunomyces, Mucor, Unclassified Fomitopsidaceae, Scedosporium, Unclassified Agaricomycetes* and *Unclassified Basidiomycota*. The observed correlations found in all kingdoms highlight distinct microbial and environmental interactions shaped by landfill activity or current operational status.

### 3.7 Co-occurrence networks of microbes in different landfill sites

Control and surface samples are often clustered together by co-occurrence network analysis, due to shared oxygen exposure and limited contact with buried refuse. However, closed and active landfill wells showed distinct microbial compositions even at similar depths, emphasizing the influence of landfill age, design, and waste input.

The differences in microbial co-occurrence networks across the site highlight key ecological shifts driven by landfill aging, substrate availability, and environmental constraints. The co-occurrence network analysis of bacterial communities in active landfill sites (Fig. 7A) revealed a highly interconnected microbial structure, with multiple clusters exhibiting strong associations. One of the most prominent clusters (green) comprises *Levilactobacillus*, *Proteiniphilum, Clostridia unclassified*, *Hydrogenispora, Caldicoprobacter*, *SBR1031_ge*, among others (e.g. OTUs 78, 84, 38, 99, and 94). These clusters are connected to another cluster (pink) composed of *Ligilactobacillus, Micromonosporaceae unclassified, Desulfallas-Sporotomaculum,* among others (OTUs 109, 74, and 43), representing lactobacilli and anaerobic bacteria involved in organic matter degradation. Another notable cluster consisted of Hydrogenispora, Halocella, and Defluvitoga (OTUs 5, 34, and 2), with Halocella being a halophilic cellulose degrader and Defluvitoga a thermophilic fermenter, suggesting adaptations to fluctuating salinity and temperature. A separate cluster featured *Hydrogenispora* and *Candidatus Caldatribacterium* (OTUs 9 and 16), along with BBMC-4 (OTU 26). *Hydrogenispora* is a known fermentative bacterium, while *Candidatus Caldatribacterium* is associated with thermophilic and anaerobic environments. Their co-occurrence suggests potential syntrophic interactions, possibly linked to production of H2. Additional clusters included *DMER64, Rhodococcus*, *Syntrophaceticus* and *Hydrogenispora* (OTUs 9, 39, 45, and 110), indicating interactions in anaerobic degradation pathways; *Paeniglutamicibacter* and *Rhabdanaerobium* (OTUs 87 and 30), which may play roles in nitrogen cycling; and *Weissella*, *Carnobacterium*, *Leuconostoc*, among others (OTUs 42, 28, 50, among others), representing fermentative bacteria associated with carbohydrate metabolism. The presence of *Thiobacillus* and *Bacillales* unclassified (OTUs 90 and 81) suggests active sulfur cycling. In active landfills, strong associations between fermenters, syntrophic bacteria, and sulfur-oxidizing taxa suggest an environment dominated by rapid organic matter degradation and microbial competition. The higher diversity and connectivity observed in bacterial networks indicate a complex and metabolically flexible community adapting to waste decomposition.

**Figure 7.**
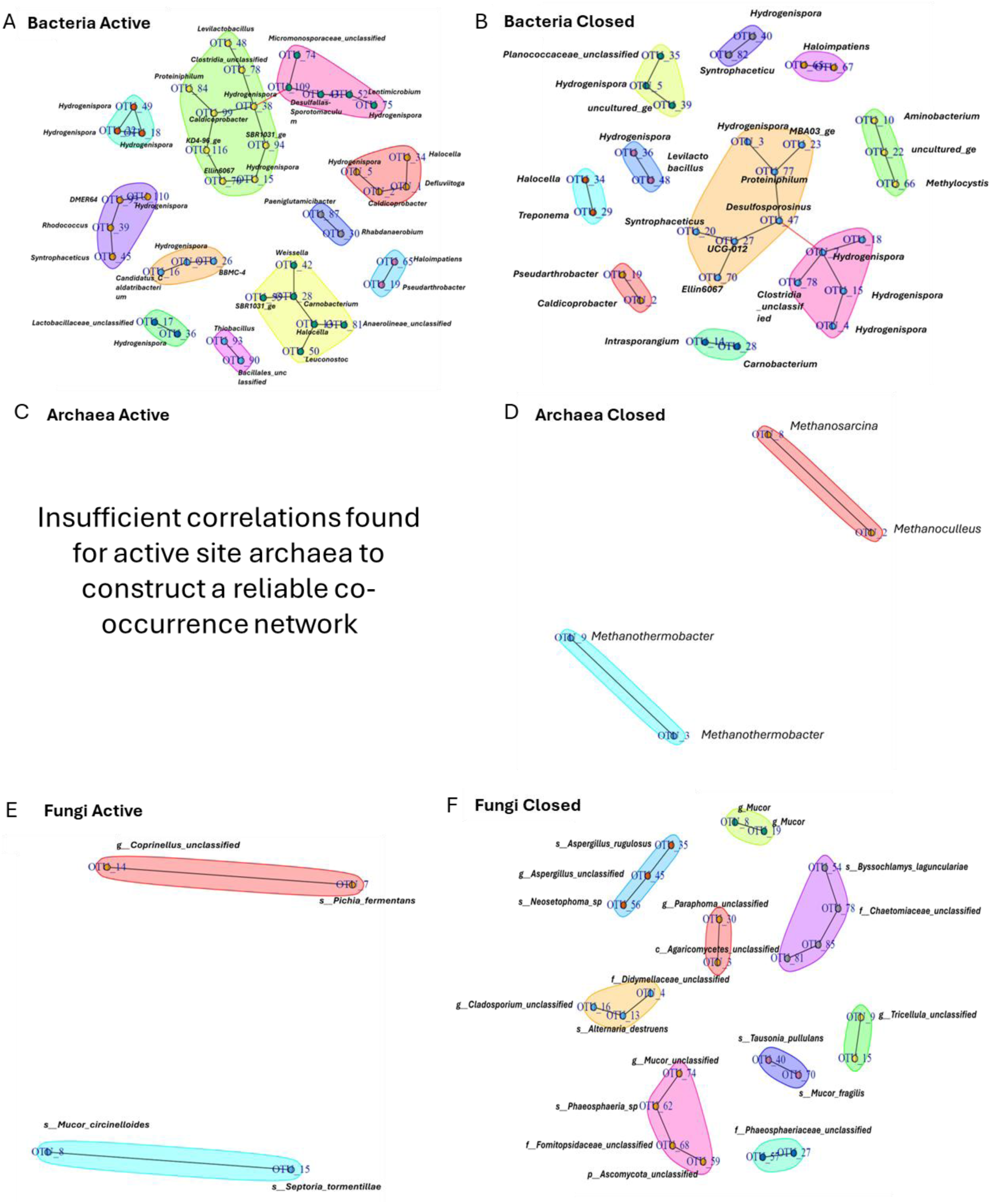
Microbial co-occurrence network analysis of microbial communities in active and closed landfill sites. (A) Bacteria Active, (B) Bacteria Closed, (C) Archaea Active, (D) Archaea Closed, (E) Fungi Active, and (F) Fungi Closed. Each network shows nodes representing operational taxonomic units (OTUs) or genera, with edges indicating significant co-occurrence relationships inferred from ensemble network analysis. Co-occurrence networks across landfills sites measured by consensus of the SPARCC and SPIEC EASI network methods. Samples are represented across landfills sites, clusters representing co-occurrence between microorganisms at the specific site, p-value: 0.03867. No network is shown for active Archaea (panel C) due to insufficient data to construct a reliable ensemble co-occurrence network.

In contrast, bacterial communities in closed landfill sites exhibited multiple well-defined clusters (Fig. 7B), suggesting structured microbial associations influenced by landfill conditions, likely driven by long-term stabilization processes. A highly connected cluster included *Desulfosporosinus, Proteiniphilum, Syntrophaceticus, UCG-012, and Ellin6067* (OTUs 20, 23, 47, 70, and 77), indicating involvement in anaerobic degradation, sulfate reduction, and syntrophic metabolism. Additionally, *Hydrogenispora* OTUs (OTUs 3, 5, 15, 18, 36, and 78) were widely distributed, suggesting a significant role in fermentation and/or/with H2 production. The strong co-occurrence of sulfate reducers (*Desulfosporosinus*) and syntrophic bacteria suggests an adaptation to anaerobic conditions where alternative electron acceptors become dominant. Other clusters included *Halocella* and *Treponema* (OTUs 34 and 29), associated with cellulose degradation, and *Planococcaceae unclassified*, *Hydrogenispora*, and an *uncultured bacterium* (OTUs 35, 39, and 5), indicating possible fermentative associations. *Pseudarthrobacter* and *Caldicoprobacter* (OTUs 2 and 19) formed another cluster, actinobacterial interactions, while *Aminobacterium, an uncultured bacterium,* and *Methylocystis* (OTUs 10, 22, and 66) suggested links to methane oxidation and nitrogen cycling.

The archaeal co-occurrence network in closed landfill sites (Fig. 7D) revealed two primary clusters: *Methanosarcina* and *Methanothermobacter*. The separation between these groups suggests niche differentiation, with *Methanosarcina* engaging in acetoclastic methanogenesis and *Methanothermobacter* relying on hydrogenotrophic methanogenesis. This differentiation may be influenced by substrate availability and redox conditions. Archaeal co-occurrence in active sites could not be plotted due to limited interactions.

Fungal co-occurrence networks in active landfill sites were less complex than those observed in closed sites, with fewer interactions among taxa (Fig. 7E). Two distinct clusters were identified in active sites, suggesting a constrained fungal community structure. One cluster included *Coprinellus unclassified* and *Pichia fermentans* (OTUs 14 and 7), indicating a possible complementary role in decomposition of organic matter such as plant waste; a second cluster consisted of *Mucor circinelloides* and *Septoria tormentillae* (OTUs 8 and 15). In contrast, fungal communities in closed landfill sites displayed multiple co-occurring clusters, suggesting structured microbial interactions (Fig. 7F). Several taxa, including *Mucor, Aspergillus, Cladosporium, Alternaria,* and *Paraphoma*, formed distinct groups, likely driven by organic matter degradation or with specific waste composition.

Other taxa, such as *Leucosporidium, Byssoclamys*, and *Piromyces*, suggested facultative/anaerobic conditions, while *Tausonia pullulans* and *Mucor fragilis* appeared to occupy specialized niches. The fungal community in closed sites also displayed greater complexity, with multiple clusters involved in organic matter breakdown, while active sites had fewer interactions, suggesting early-stage fungal colonization.

## 4. Discussion

Analysis of the landfill ecosystem revealed two simultaneous ecological processes: microbial community succession and niche differentiation across landfills ages and sites. Succession was observed across time (depth), reflecting the transition of microbial communities as environmental conditions evolved with time, such as oxygen depletion, waste burial, and decomposition. We posit that landfill phases can be predicted from microbial community composition and our observed pattern of succession, along with the characteristic metabolisms within each phase such as aerobic degradation, fermentation, and methanogenesis. Niche differentiation was simultaneously observed at every depth within the landfill, where diverse microenvironments stemming from specific waste inputs allowed distinct taxa to persist despite the overall microbial community succession pattern.

### 4.1 Biogeochemical profiles show clear succession of phase-linked transitions driven by depth, age and oxygen depletion

The most abundant microbial taxa at each depth reflect potential dominant metabolic functions and therefore the following landfill biological phases of aerobic degradation, anaerobic acid production (fermentation), and methanogenesis. Previous studies support this pattern, with surface layers enriched in oxygen-tolerant taxa and deeper dominated by anaerobic communities associated with fermentation and methanogenesis (S. jia Liu et al., 2019; Y. X. Wang et al., 2025). Here, we extend this concept by interpreting vertical zones as functional analogs to temporal decomposition phases, offering a spatial framework for landfill functional succession.

Surface layers were enriched with aerobic degraders likely adapted to and inoculated by fresh refuse inputs. At mid-depths, the microbial community shifted to being dominated by fermenters that produce intermediate metabolites such as hydrogen and short-chain fatty acids. In deeper layers, methanogens that utilize these fermentation byproducts for methane production became the primary taxa. These transitions are driven by the thermodynamic favorability of metabolic pathways under different environmental conditions, including electron acceptor availability (oxygen, nitrate, sulfate) and carbon source composition (e.g., food waste, plastics, lignocellulose). This pattern reflects secondary establishment processes, where initial colonizers modify the environment, such as depleting oxygen and producing metabolites, which in turn support the growth of fermenters and methanogens. The shifts in microbial community composition are also a factor of time, or depth, since the deeper layers of the landfill are older in age than the surface sites. This sequential pattern is characteristic of microbial succession and has been widely observed in landfill ecosystems (Dong et al., 2015). Compared to other landfills, the Dane County Landfill exhibited typical successional stages but also site-specific microbial patterns, likely shaped by factors such as waste input, operational practices, and local climate.

The biogeochemistry observed in the surface and top layers of the active sites of the landfill was consistent with Phase 1 processes, i.e., decomposing organic waste by aerobic bacteria, consuming oxygen and breaking down long molecular chains in organic matter. Here, we observed predominance of aerobic bacteria and fungi in the surface and shallower active site samples, which are known to initiate early waste degradation. Fungi play a crucial role in the early stages of organic waste degradation by breaking down complex compounds (carbohydrates, lipids and proteins), into lower molecular weight substances such as sugars, fatty acids, and amino acids. This process primarily results in the production of CO₂ and H₂O, along with the formation of stabilized humic substances that can be further utilized by other microorganisms. Our results are supported by previous studies showing that aerobic bacteria and fungi predominate in top landfill layers due to O2 availability (Sekhohola-Dlamini & Tekere, 2020; Uz et al., 2003; Y. nan Wang et al., 2021). These surface communities, with minimal exposure to buried waste, were more similar in composition to the relatively undisturbed soil control samples from the fill dirt, with changes in the landfill surface samples likely representing early-stage microbial assemblages.

As O2 is depleted, the transition to the anaerobic acid phase (Grégoire et al., 2023) began in the two active site wells around 30-40 feet in depth and by 10 feet in the closed site wells at the time of our sampling. Depths we assigned to this phase were characterized by the increase of fermentative bacteria, and a decrease in the relativeabundance of the fungal community dominated by anaerobic or stress-tolerant fungi, consistent with conditions favoring anaerobic decomposition (Sekhohola-Dlamini & Tekere, 2020). In the closed sites, we observed an increased abundance of anaerobic bacteria and archaea across all depths relative to active site wells. Within each well, archaea increased in abundance with depth, indicating the transition to the methanogenesis phase in depths from 40-70 ft in active sites and across all depths of closed sites. In these depths, methanogens such as *Methanosarcina*, *Methanobacterium*, and *Methanoculleus* became dominant, signaled by active methane production. This is further supported by the co-occurrence network analysis which revealed strong associations between syntrophic fermenters and methanogens, particularly in closed sites, consistent with mature anaerobic systems producing methane (Barlaz et al., 1990; Grégoire et al., 2023).

The microbial community patterns were also reflected in the trends in soil color. Surface and control (fill-dirt) samples displayed similar red or clay tones, likely due to limited exposure to waste and aerobic conditions. In active landfill sites, color gradients with depth can reflect a shift from aerobic to anaerobic conditions, degradation of waste, or waste input variations (Long et al., 2016; P. Wang et al., 2022). Closed landfill sites exhibited consistent brown or darker tones across all depths, which could indicate more advanced decomposition or the fact that the landfill being capped limits oxygen and enhances stabilization. These differences highlight the roles of landfill physical properties and age in shaping soil characteristics. Together, these biotic and abiotic findings present a high-resolution picture of microbial succession, linking changes in community composition with depth, oxygen availability, waste exposure, metabolic function, and geochemistry, ultimately having implications for methane production and waste stabilization.

### 4.2. Niche Differentiation: Microcompartments and Preservation of Community Richness

Succession describes temporal transitions in microbial communities. This was observed in the Dane County Landfill by percentages of fungi, bacteria and archaea, and was aligned with known gas production phase transitions for municipal sanitary landfills. Niche differentiation refers to the spatial and resource variation of microbial communities at individual landfill depths and locations. Our data indicate that niche differentiation was strongly influenced by the heterogeneous composition of refuse which generates local physicochemical gradients and therefore distinct microhabitats.

#### 4.2.1 Evidence of Microcompartment Formation

Refuse at any given depth is a heterogenous mixture composed of food waste, plastics, wood, paper, construction debris, and more. Our physicochemical data suggests the formation of microcompartments within every depth driven by the particular item of trash present at that location. Firstly, we found that the presence of trash changes the physicochemical parameters as seen in the significantly lower concentrations of ions, metals, OM, and TN in comparison to control fill dirt samples that have never been exposed to trash compared to the trash-exposed samples. Secondly, for trash-exposed samples, there were no clear trends by depth for any of the physicochemical parameters which suggests that these values are determined by the types of materials at that depth.

Geochemical data showed localized spikes in ammonia, sulfate, and heavy metals, evidence of environmental patchiness that facilitated the coexistence of niche-adapted taxa (Fig. 5). For example, at depths such as 90 ft in the closed site wells, the high abundance of taxa such as *Psychrobacillus* was strongly correlated with high levels of copper at only two locations that were sampled. This is consistent with previous studies; the age of waste appears to significantly influence these parameters (Song et al., 2015). The niche differentiation pattern, in both active and closed wells, suggests that environmental conditions at greater depths may not only inhibit growth of unadapted microbes but also enhance waste degradation by taxa specialized to survive those conditions (Long et al., 2016; P. Wang et al., 2022). This heterogeneity in physicochemical parameters in a landfill led to the formation of distinct microhabitats, each supporting unique microbial populations.

##### 4.2.1.1 Bacteria

In surface and active sites, the dominance of *Proteobacteria, Actinobacteria*, and *Firmicutes* aligns with previous landfill studies (Ke et al., 2022; Li et al., 2022; S. jia Liu et al., 2019; Morita et al., 2020; Sawamura et al., 2010; Schupp et al., 2020; P. Wang et al., 2022; Y.-N. Wang et al., 2021; Xu et al., 2017). The phylum-level similarity between control and surface samples may be due to the small concentration of trash relative to soil in landfill surface sites (Fig.4).

At greater depths (10 ft and below) in active sites, there is a marked increase in *Firmicutes* and a decline in *Proteobacteria* (Fig.4). This shift is consistent with the anaerobic conditions deeper in the landfill, where *Firmicutes* play a key role in degradation (S. jia Liu et al., 2019). *Firmicutes* are among the predominant phyla in landfill environments, facilitating the breakdown of organic matter under oxygen-depleted conditions. However, contrary to earlier reports, we observed an increase in *Firmicutes* at deeper portions of the active site and in the closed sites of the Dane County Landfill (Y.-N. Wang et al., 2021). This discrepancy may be due to landfill-specific conditions including continuous waste input, temperature fluctuations, and the lack of capping in active sites. In closed sites, evidence of microcompartments and niche differentiation included predominant presence of *Hydrogenispora, Caldicoprobacter, and Defluviitoga*, which thrive in the warm, anaerobic conditions that are found stably in these locations. Likewise, the presence of unique taxa at the deepest depths sampled, which are influenced by the proximity to the plastic liners and leachate collection system components, further suggest that distinct environments within the landfill create niches that select for certain taxa.

The bacterial community and physicochemical parameters of the leachate samples were distinct from the landfill or control samples, reflecting that the leachate is a unique and fluctuating environment. Click or tap here to enter text. Research conducted across 16 leachate landfills in the U.S. revealed that the dominant bacterial phyla were *Proteobacteria*, *Firmicutes*, and *Bacteroidetes (*Stamps et al., 2016), but each leachate sample harbored distinct microbial communities, with only 2.9% of detected OTUs shared across all landfills. This variation is likely influenced by factors such as chemical composition, waste age, and metal concentrations. Similar findings were reported in a study of 12 landfill leachates in China, which also highlighted site-specific microbial diversity (Zhao et al., 2021). Our results aligned with these previous studies, as *Proteobacteria* and *Firmicutes* were the dominant bacterial phyla in our leachate samples. However, we did not observe the presence of obligately anaerobic *Bacteroidetes*, possibly due to the open leachate facilities at the DCL. Additionally, the unique taxa identified in our leachate samples possess key metabolic capabilities, including heavy metal removal, dye and hydrocarbon degradation, nitroaromatic compound breakdown, polymer decomposition, and ammonia removal (Xiao et al., 2024).

##### 4.2.1.2 Archaea

Control samples showed a dominant presence of specific genera, while surface layers were characterized by *Nitrososphaeraceae_ge* (Otu00018), *Candidatus Nitrosocosmicus* (Otu00016), *Methanosarcina* (Otu00008), *Methanobacterium* (Otu00007), and *Candidatus Methanoperedens* (Otu00020). The nitrifying taxa identified in the surface layers are common soil archaea and belonging to the group of ammonia-oxidizing archaea (AOA), which play a critical role in the nitrogen cycle. These archaea have also been identified in deep oxidation ditches of wastewater treatment plants that process landfill leachate (Y. Yang et al., 2021). As depth increases, a shift toward methanogenic taxa, such as, *Methanobacterium and Methanosarcina* is observed in active sites. While some depths exhibit varying levels of abundance, these taxa consistently dominate across different layers. Their presence has been widely reported in landfills of various ages and depths (Dong et al., 2015), and crucial roles in methane production, with *Methanomicrobiales* and *Methanosarcinales* being associated with anaerobic methane oxidation (ANME) and reverse methanogenesis (Dong et al., 2015).

The unique archaeal composition in our leachate samples, particularly the presence of *Methanosaeta-* a strict acetoclastic methanogen that exclusively utilizes acetate as a substrate, along with *Candidatus Nitrosocosmicus*, highlights the dual role of these taxa in both methanogenesis and nitrogen cycling under contaminant-rich conditions. The dominant archaeal taxa also include *Methanocorpusculum-* a hydrogenotrophic methanogen that produces methane by reducing CO₂ with H₂, and *Candidatus Methanoplasma,* , a methyl-reducer that uses H₂ as an electron donor. The high abundance of putative hydrogenotrophic *Methanomicrobiales* in leachate is likely linked to the presence of syntrophic bacteria that metabolize fatty acids and generate H₂, facilitating interspecies hydrogen transfer (Staley et al., 2012). The presence of these archaea in landfill leachate is consistent with recent studies that highlight their role in methanogenic processes (Co & Hug, 2021; Grégoire et al., 2023; Sauk & Hug, 2022) and their importance in landfill biogeochemical processes.

##### 4.2.1.3 Fungi

The phylum *Ascomycota* dominated across all landfill sites that align with previous findings, highlighting their functional role in organic matter degradation (Sekhohola-Dlamini & Tekere, 2020; Szulc et al., 2022; Ye et al., 2020, 2024). The composition of specific fungal species varied across the landfill. In the surface layers of active landfill sites, environmental conditions such as higher oxygen levels and frequent substrate inputs appeared to favor a narrower range of taxa, resulting in the dominance of specific fungi/yeasts such as *Penicillium, Pichia fermentans*, and *Teunomyces* (OTU00012) which are associated with food degradation and have also been reported in other landfill environments (G. Fu et al., 2022; Shrivastava et al., 2023; Szulc et al., 2022; Wolski, 2023).

In contrast, closed landfill sites had fungal communities characterized by taxa with specialized adaptations to persist in more stable, anoxic, and nutrient-depleted microhabitats. Species such *Pseudeurotium_unclassified* (OTU00005), *Scedosporium dehoogii* (OTU00002), *Pseudallescheria boydii* (OTU00001), *Mucor circinelloides* (OTU00008), and *Didymellaceae_unclassified* (OTU00004) were prevalent, consistent with their ability to tolerate or even thrive under anaerobic conditions (Munir et al., 2024; Yadav et al., 2023). The unique dominance of certain taxa (e.g., Cladosporium in sample 548) further suggests fine-scale niche differentiation potentially influenced by interactions with landfill infrastructure such as plastics liners and its relation with landfill systems. (G. Fu et al., 2022; Mondal et al., 2023; Szulc, Okrasa, Nowak, et al., 2022).

Overall, leachate samples presented a unique fungal profile, dominated by *Dipodascaceae_unclassified* and *Yarrowia lipolytica*. The specialized metabolic capabilities of these fungi/yeast to live in polluted environments, enable them to exploit ecological niches that are inaccessible to less metabolic diverse organisms (Zinjarde et al., 2014). Lower overall diversity in leachate samples, consistent with prior studies, reflects strong environmental filtering that restricts community assembly to functionally specialized taxa (Ye et al., 2020).

#### 4.2.2 High Microbial Richness and Uneven Community Structure

Our findings did not exhibit a decrease in diversity with age or depth, despite microbial diversity tending to decrease at higher temperatures (Schupp et al., 2020). Beta and gamma diversity were high, indicating the coexistence of diverse communities across landfill zones. Alpha-diversity patterns, likewise, varied but remained high across age and depths and sites. These results mirror diversity patterns reported in other landfills (Liu et al., 2019; Sawamura et al., 2010; P. Wang et al., 2022; Ye et al., 2020). Control and surface samples showed the highest Shannon diversity indices for all microbial communities, a finding that aligns with previous studies on bacterial communities. For instance, Sawamura et al. (2010) investigated landfills in Japan using real-time PCR assays and observed lower Shannon–Weaver diversity indices at 3.0 m and 11.5 m depths. They reported variation of archaeal and bacterial copy numbers, which may explain similar stratified patterns observed in our current landfill study. Similarly, other studies exploring fungal diversity in refuse samples followed comparable trends as ours, with Shannon diversity indices ranging from 2.0 to 3.5 (Ye et al., 2024a).

Our findings support the concept of niche differentiation, since alpha diversity is a function of both species’ richness and evenness. Species richness was high throughout, with thousands of individual taxa found in every sample. This persistence of species richness is not surprising, due to the heterogeneity of waste, microbial metabolisms and other components that constitute landfills. Despite this richness in diversity, community evenness was low, suggesting that certain taxa thrive under specific environmental constraints. Venn diagrams confirmed limited OTU overlap across sites and depths, with only 383 bacterial OTUs, 19 archaeal OTUs, and 178 fungal OTUs shared across all samples. This highlights the presence of highly specialized and site-specific microbial assemblages shaped by environmental gradients (Staley et al., 2012). This is reflective of broader ecological principles observed in other landfills environments, where beta diversity remains high due to localized selection pressures and habitat differentiation (Liu et al., 2019; Sawamura et al., 2010; P. Wang et al., 2022; Ye et al., 2020). Niche differentiation in the landfill thus plays a central role in preserving microbial richness and functional diversity despite environmental stress. Altogether, these findings underscore that microbial diversity in landfills is not solely the result of vertical succession, but also of horizontal variability and microhabitat specialization, key factors that must be considered when studying biogeochemical cycling and planning landfill management strategies.

### 4.3. Implications for Landfill Management

The findings from this study have implications for landfill management strategies, especially in relation to waste stabilization, methane recovery, and bioremediation potential. By understanding how microbial communities change with landfill age, depth, and geochemistry, managers can better predict decomposition stages and optimize operational decisions. One key implication involves gas capture and renewable energy generation (RGE). As our study confirmed, methane production is closely associated with deeper layers and older waste in both active and closed sites. The presence of syntrophic fermenters and hydrogenotrophic methanogens such as *Hydrogenispora* and *Methanothermobacter* suggests that enhancing these microbial interactions, potentially by modulating temperature or substrate availability, could improve methane yields. This could inform strategies like targeted heating, bioaugmentation, or pH balancing to enhance biogas recovery while minimizing greenhouse gas emissions (Shao et al., 2023). Beyond management implications, results presented here are of additional importance because Artesols have recently been proposed as a new soil order that is characterized by a single, dominant soil forming factor (human activity). An initial characterization of this Artesol type will serve to begin to develop our understanding of the microbiome of this new soil order.

Our results also reveal that leachate microbial communities are adapted to extreme conditions, including elevated metal concentrations, salinity, and ammonia. Dominant genera such as *Exiguobacterium*, *Pseudomonas*, and *Yarrowia lipolytica* show potential for bioremediation of contaminants and plastic additives, reinforcing the value of monitoring and managing leachate not only as a waste product but also as a microbial ecosystem with functional potential (Zhao et al., 2021; Xiao et al., 2024; Zinjarde et al., 2014).

Our *in situ* landfill sampling results, in conjunction with previous field-scale landfill test cells and smaller bioreactor studies (Barlaz et al., 1997; Giangeri et al., 2022; Staley et al., 2012), point to implications for gas production and control. Previous studies highlighted the importance of understanding landfill types like anaerobic, leachate recirculated anaerobic, semi-aerobic and aerobic, and the use of leachate recirculation for the quicker stabilization of landfills (Sekman et al., 2019). Likewise, laboratory-scale reactors and bioreactors have been used to investigate landfill microbiology and characterize its process of biodegradation in a controlled environment. For instance, previous research using reactors identified that refuse composition, for example increasing carbohydrates like cellulose and hemicellulose, increases its production of methane (Barlaz et al., 1997). Consistent with these findings, our study shows that microbial community succession in landfills unfolds over a decadal scale before reaching the stage of rapid methanogenesis. This long timescale provides context for the practice of leachate recirculation, which can shorten stabilization periods by accelerating the establishment of methanogenic communities, while not significantly altering their composition (Barlaz et al., 1990).

While leachate recirculation can enhance waste stabilization and methane recovery, potential trade-offs also need to be considered. In particular, landfills with construction and demolition waste or other sources of sulfate, a stabilized anaerobic community may also produce higher amounts of gas contaminants like H₂S, which can interfere with methane recovery systems and pose hazards at collection facilities. Monitoring sulfate-reducing bacteria and optimizing conditions that favor methane over H₂S production may mitigate these risks. (Q. Xu et al., 2014; Q. Xu & Townsend, 2014)

Finally, the observed microbial heterogeneity emphasizes the value of stratified sampling and tailored management at different depths and zones. A one-size-fits-all approach may overlook the dynamic interactions shaping microbial succession and niche partitioning. Instead, adaptive strategies that account for microbial phase transitions and diversity patterns can enhance both environmental outcomes and resource recovery in landfill operations. Therefore, incorporating knowledge of microbial ecology into landfill management could improve waste treatment efficiency, reduce environmental contamination, and inform the development of biotechnologies that harness landfill microbial communities for energy and remediation purposes.

## 5. Conclusion

In this study, we sampled a landfill across depths and ages in Madison, Wisconsin to understand how landfill history and management strategies impact microbial community structure and function. Microbial diversity patterns revealed stratification across depths, with surface and control samples exhibiting the highest richness. Active sites harbored metabolically diverse bacterial communities, while closed sites demonstrated microbial stabilization, favoring anaerobic taxa involved in methanogenesis and organic matter breakdown. Fungal diversity was site-specific, with aerobic fungi dominating surface layers and anaerobic, stress-tolerant species thriving in deeper refuse. Leachate samples exhibited distinct microbial communities adapted to leachate conditions, with bacterial and fungal taxa specialized in contaminant degradation.

Co-occurrence network analysis underscored the ecological shifts between active and closed sites, with active landfills supporting dynamic microbial interactions associated with organic matter fermentation, while closed landfills exhibited structured networks favoring methane-producing archaea and syntrophic bacteria. The observed persistence of methanogenic taxa across depths and landfill ages indicates that methane production remains a dominant microbial function, regardless of landfill management practices.

Overall, our findings contribute to a deeper understanding of landfill microbial ecology, emphasizing the role of physicochemical gradients in shaping microbial communities, which are ultimately shaped and can also, can inform, human management choices. In coming decades, landfill management strategies such as composting and waste diversion will be more implemented (Tominac et al., 2021). As we move towards a goal of methane mitigation and pollution reduction, understanding the long-term impacts of microbial succession should illuminate the remarkable bioremediation potential of landfill microbial communities.

Funding information

This work supported by the following fundings: NSF Graduate Research Fellowship Program (GRFP) #2137424, Science and Medicine Graduate Research Scholars (SciMed GRS) and Biotechnology Training Program (BTP) at the University of Wisconsin–Madison T32 GM135066.

## Acknowledgements

We would like to acknowledge the Dane County Landfill (DCL) for providing us with the opportunity to collect samples during the installation of methane gas wells. We are especially grateful to the DCL staff—Allison Rathsack (Lead Project Engineer), John Welch (Director), and Jack Hansen (Intern)—for granting access to the landfill and offering invaluable guidance during sampling. We extend our thanks to Xiadong Wang from the geoengineering lab at UW-Madison for lending us essential equipment for the landfill sampling. Special thanks to Geoffrey Siemering and Francisco Arriaga from the UW-Madison Soils Department for allowing us to borrow the X-ray fluorescence machine (Innov-X Systems), Thermo Scientific Orion Star™ electrode, crucibles, and muffle furnace, as well as for their guidance on sample preparation and analysis. We would also like to thank James Lazarcik from the Water Science and Engineering Laboratory at UW-Madison for his training and support with ion chromatography (IC) and sample preparation for anion and cation analysis.

## Author contributions

**D.R.R.:** Conceptualization, Methodology, Validation, Formal Analysis, Investigation, Writing-Original Draft & Editing, Data Processing and Visualization, NSF-GRFP Funding, **L.H.:** Conceptualization, Methodology – sample collection, qPCR and MiSeq library preparation and sequencing. **W.C.:** Data Processing and Visualization, Writing - Sequence Analysis and Statistical analysis. **P.Q. T.**: Data Processing of Microbiome data, Formal Analysis, Visualization, Writing – Review and Editing. **H.N.F.**- Methodology – Sample preparation for Ion Chromatography. **F.J.S**.: Methodology – Sample Collection. **D.A.R.K:** Methodology – Sample Collections. **J.S.:** Methodology: Mentored during MiSeq library preparation and provided protocols of data processing (Mothur). **G.C.** Methodology – sequence analysis. **Z. F.:** G.C. PI, Methodology – sequence analysis, Writing – Review and Editing **G. S.:** J.S PI. **B.G.F**: Mentoring, Writing – Review and Editing.**, A.R.:** DCL Staff**, E.L.W. M**.: Conceptualization, Resources, Project Administration, Supervision, Funding Acquisition, Writing-Review & Editing.

