## Supplemental Figures and Tables for "Patterns of Microbial Succession and Niche Differentiation Across Depth and Age in a Landfill have Implications for Management Strategies"

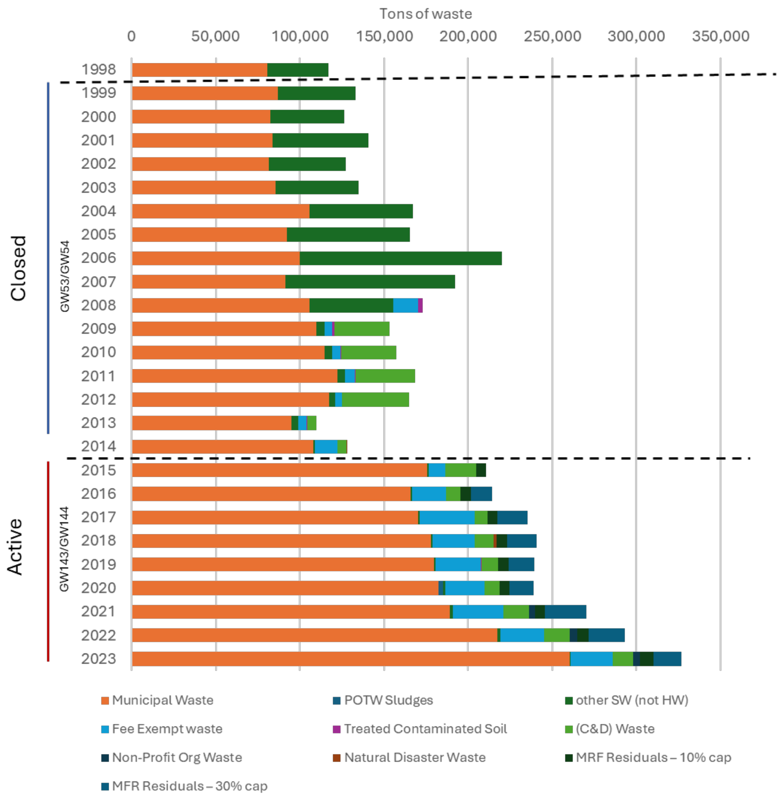


Figure S1: Trends in Waste Deposition at the Landfill Over Time (1998–2023). Stacked bar chart illustrates the total waste deposited in the DCL landfill annually, categorized by waste type. Data sourced from the Wisconsin Department of Natural Resources. Solid Waste Tip Fees and Landfill Tonnage Reports. Retrieved from https://dnr.wisconsin.gov/topic/Landfills/Fees.html

Table S1: Overall waste in tons and their categories through the years

| Year | Municipal Waste | POTW Sludges | other SW (not HW) | Fee Exempt waste | Treated Contaminated Soil | (C&D) Waste | Non-Profit Org Waste | Natural Disaster Waste | MRF Residuals – 10% cap | MFR Residuals – 30% cap |
|---|---|---|---|---|---|---|---|---|---|---|
| 1998 | 80,506 | 0 | 36,272 | 0 | 0 | n/a | n/a | n/a | n/a | n/a |
| 1999 | 86,958 | 0 | 46,174 | 0 | 0 | n/a | n/a | n/a | n/a | n/a |
| 2000 | 82,317 | 0 | 44,238 | 0 | 0 | n/a | n/a | n/a | n/a | n/a |
| 2001 | 83,661 | 0 | 57,133 | 0 | 0 | n/a | n/a | n/a | n/a | n/a |
| 2002 | 81,676 | 0 | 45,669 | 0 | 0 | n/a | n/a | n/a | n/a | n/a |
| 2003 | 85,620 | 0 | 49,389 | 0 | 0 | n/a | n/a | n/a | n/a | n/a |
| 2004 | 105,738 | 0 | 61,316 | 0 | 0 | n/a | n/a | n/a | n/a | n/a |
| 2005 | 92,254 | 0 | 72,890 | 0 | 0 | n/a | n/a | n/a | n/a | n/a |
| 2006 | 99,763 | 0 | 120,125 | 0 | 0 | n/a | n/a | n/a | n/a | n/a |
| 2007 | 91,592 | 0 | 100,576 | 0 | 0 | n/a | n/a | n/a | n/a | n/a |
| 2008 | 105,871 | 0 | 49,737 | 14,778 | 2,644 | n/a | n/a | n/a | n/a | n/a |
| 2009 | 109,806 | 0 | 4,671 | 4,671 | 1,426 | 32,644 | 0 | n/a | n/a | n/a |
| 2010 | 114,518 | 0 | 4,780 | 4,780 | 509 | 32,756 | 0 | n/a | n/a | n/a |
| 2011 | 122,370 | 0 | 4,272 | 6,107 | 474 | 35,429 | 0 | 0 | n/a | n/a |
| 2012 | 117,259 | 12 | 3,789 | 3,945 | 160 | 39,751 | 0 | 0 | n/a | n/a |
| 2013 | 94,906 | 2 | 3,916 | 5,082 | 494 | 5,198 | 0 | 0 | n/a | n/a |
| 2014 | 108,071 | 0 | 1,012 | 13,197 | 0 | 5,561 | 0 | 275 | n/a | n/a |
| 2015 | 175,500 | 0 | 1,082 | 9,727 | 152 | 18,122 | 0 | 0 | 6,127 | 0 |
| 2016 | 165,811 | 0 | 1,045 | 20,187 | 0 | 8,434 | 0 | 0 | 6,182 | 12,466 |
| 2017 | 170,141 | 0 | 909 | 32,853 | 0 | 7,595 | 0 | 0 | 5,904 | 17,787 |
| 2018 | 177,816 | 0 | 906 | 25,155 | 0 | 11,065 | 0 | 2,129 | 5,904 | 17,787 |
| 2019 | 179,688 | 0 | 770 | 27,034 | 348 | 10,143 | 0 | 0 | 6,086 | 15,242 |
| 2020 | 182,532 | 2,595 | 1,206 | 23,507 | 0 | 8,922 | 0 | 0 | 5,897 | 13,997 |
| 2021 | 189,067 | 0 | 1,674 | 30,098 | 0 | 15,352 | 3,759 | 0 | 5,786 | 24,737 |
| 2022 | 217,536 | 0 | 1,740 | 26,015 | 0 | 15,096 | 4,421 | 0 | 6,841 | 21,584 |
| 2023 | 260,531 | 0 | 402 | 24,867 | 0 | 12,025 | 4,200 | 0 | 8,226 | 16,602 |

Data sourced from the Wisconsin Department of Natural Resources. Solid Waste Tip Fees and Landfill Tonnage Reports. Retrieved from https://dnr.wisconsin.gov/topic/Landfills/Fees.html

Table S2: RT-qPCR Primers used for bacteria, archaea and fungi

| | Region | Primer Set | Sequence | Tm° (annealing temperature of primers) | Amplicon base (bp) | Reference |
|---|---|---|---|---|---|---|
| 16S rRNA Bacteria | V4 | 341F | CCT AYG GGR BGC ASC AG | 64.6 | 466 | (Takahashi et al., 2014) |
| | | 806R | GGA CTA CNN GGG TAT CTA AT | 55.2 | | |
| 16S rRNA Archaea | V6-V8 | arch349F | GYG CAS CAG KCG MGA AW | 76.9 | 422 | (Takahashi et al., 2014) |
| | | arch806R | GGA CTA CVS GGG TAT CTA AT | 55.1 | | |
| ITS Fungi | ITS1 | ITS1f | TCC GTA GGT GAA CCT GCG G | 61.5 | 267 | (Q. Fu et al., 2020; Takahashi et al., 2014) |
| | | 5.8 s | CGC TGC GTT CTT CAT CG | - | | |

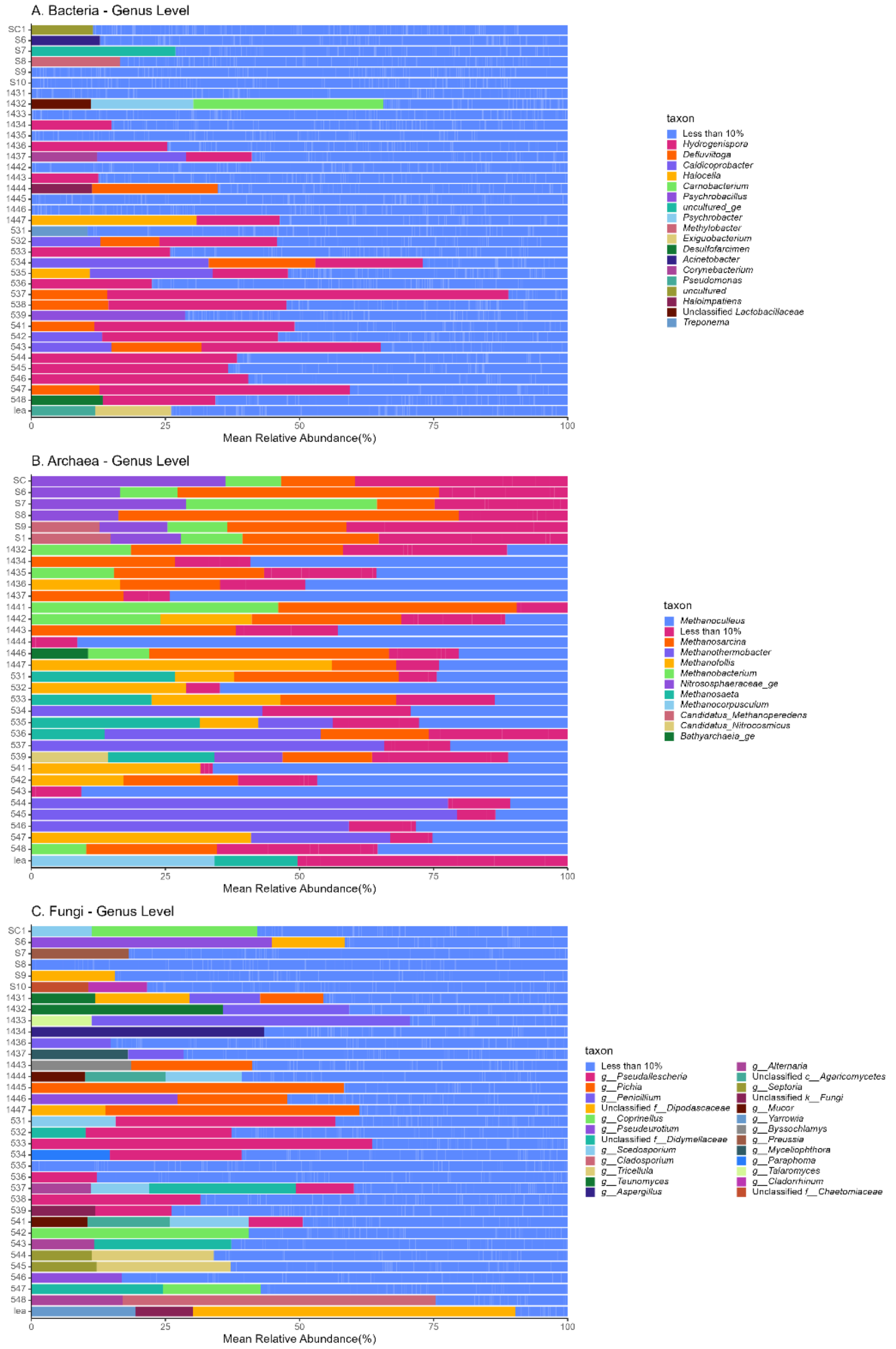


Figure S2: Relative abundance bacterial (A), archaeal (B), and fungal (C) genera across landfill samples. Stacked bar plots show the genus-level composition of microbial communities in each sample, highlighting genera that individually comprise less than 10% of the total community genera. The category “Less than 10%” aggregates all low-abundance genera not individually labeled in the legend. Full OTU tables can be found in the public KBase narrative.

Table S3: Summary of Total Numbers of Taxonomic Diversity and OTU Counts

| Kingdom | Phylum | Class | Order | Family | Genus | Species | Total OTU counts |
|---|---|---|---|---|---|---|---|
| Bacteria | 46 | 142 | 319 | 541 | 1084 | N/A | 4175 |
| Archaea | 7 | 13 | 15 | 29 | 42 | N/A | 214 |
| Fungi | 18 | 66 | 136 | 318 | 636 | 1007 | 2785 |

Table S4: Physicochemical parameters and environmental factors across samples

| sample_id | NO3- | NO2- | SO42- | PO43- | Na+ | Ca2+ | Mg2+ | K+ | NH4+ | Ag | As | Cr | Cu | Fe | K | Mn | Mo | Nb | Ni | Pb | Rb | Sn | Sr | Ti | V | Y | Zn | Zr | TN | OM | EC | pH | Temperature | Depth | Well Name |
|---|---|---|---|---|---|---|---|---|---|---|---|---|---|---|---|---|---|---|---|---|---|---|---|---|---|---|---|---|---|---|---|---|---|---|---|
| 1431 | NA | NA | 1463.6 | NA | 3355.2 | 6382.1 | 826.1 | 1227.6 | 491.6 | NA | NA | 29.0 | 56.0 | 5146.5 | 8027.0 | 218.0 | NA | 3.4 | NA | 324.0 | 20.5 | NA | 237.0 | 1136.5 | 30.0 | 5.8 | 758.5 | 110.0 | 10679.0 | 40.1 | 6.7 | 6.2 | 76 | 10 | GW143 |
| 1432 | NA | NA | 1510.5 | NA | 1663.9 | 4832.1 | 1030.9 | 791.6 | NA | NA | NA | 30.0 | 63.0 | 5106.5 | 7258.0 | 321.0 | NA | NA | NA | 389.0 | 18.2 | NA | 397.0 | 1385.0 | 29.0 | 7.3 | 362.0 | 96.0 | 6786.2 | 33.7 | 5.4 | 6.9 | 42 | 20 | GW143 |
| 1433 | NA | NA | 1509.7 | NA | 3209.0 | 5376.7 | 777.1 | 2130.6 | NA | NA | NA | 39.0 | 105.0 | 8528.0 | 8386.0 | 319.0 | NA | 5.3 | 15.0 | 465.0 | 23.3 | NA | 245.0 | 1462.5 | 38.0 | 8.5 | 766.5 | 123.0 | 8859.3 | 41.5 | 6.9 | 6.2 | 87 | 30 | GW143 |
| 1434 | NA | NA | 269.2 | NA | 833.4 | 262.6 | 95.8 | 497.9 | 272.3 | NA | 15.0 | 77.0 | 71.0 | 17900.5 | 10426.0 | 442.0 | NA | 6.4 | 42.0 | 172.0 | 26.4 | NA | 242.0 | 1986.0 | 38.0 | 9.5 | 552.5 | 184.0 | 5237.3 | 16.0 | 1.7 | 7.5 | 92 | 40 | GW143 |
| 1435 | NA | NA | 535.6 | NA | 684.7 | 642.8 | 154.6 | 71.8 | 241.8 | 34.0 | 10.0 | 62.0 | 182.0 | 51596.0 | 11979.0 | 823.0 | 4.8 | 5.9 | 30.0 | 108.0 | 35.5 | 158.0 | 172.0 | 1797.5 | 22.0 | 14.1 | 289.5 | 204.0 | 3122.5 | 12.7 | 1.6 | 7.9 | 102 | 50 | GW143 |
| 1436 | NA | NA | 702.2 | NA | 2044.5 | 947.1 | 147.0 | 294.0 | 493.2 | NA | NA | 43.0 | 136.0 | 18230.5 | 11471.0 | 1171.0 | NA | 10.4 | 29.0 | 272.0 | 37.1 | NA | 168.0 | 1934.0 | 42.0 | 12.3 | 366.5 | 424.0 | 4508.8 | 16.4 | 2.4 | 7.1 | 100 | 60 | GW143 |
| 1437 | NA | NA | 1588.8 | NA | 1970.7 | 2704.7 | 168.5 | 529.0 | 282.2 | NA | NA | 66.0 | 88.0 | 15451.0 | 10126.0 | 439.0 | NA | 6.1 | 28.0 | 113.0 | 31.6 | NA | 231.0 | 1637.0 | 32.0 | 10.1 | 704.0 | 223.0 | 4401.8 | 22.7 | 4.3 | 7.1 | 110 | 70 | GW143 |
| 1441 | NA | NA | 1557.1 | 9.4 | 1668.8 | 8510.9 | 528.0 | 446.6 | 100.0 | NA | 14.0 | 34.0 | 123.0 | 7385.5 | 8367.0 | 595.0 | NA | 4.9 | 23.0 | 241.0 | 23.3 | NA | 449.0 | 1573.5 | 32.0 | 10.6 | 479.5 | 155.0 | 6027.8 | 32.2 | 4.9 | 6.6 | 90 | 10 | GW144 |
| 1442 | NA | 20.9 | 1792.4 | NA | 2294.5 | 7993.0 | 759.7 | 339.8 | 201.6 | NA | NA | 27.0 | 102.0 | 6550.5 | 5358.0 | 250.0 | NA | 4.3 | NA | 262.0 | 17.5 | NA | 537.0 | 2182.5 | 23.0 | 5.7 | 327.5 | 759.0 | 5477.9 | 29.6 | 5.5 | 7.0 | 74.5 | 20 | GW144 |
| 1443 | NA | NA | 883.5 | NA | 659.2 | 3293.7 | 315.0 | 130.6 | 60.6 | NA | NA | 35.0 | 45.0 | 13273.0 | 12412.0 | 586.0 | NA | 6.8 | 21.0 | 48.0 | 31.9 | NA | 188.0 | 1701.0 | 36.0 | 11.3 | 217.0 | 207.0 | 3549.6 | 9.3 | 2.7 | 7.5 | 80 | 30 | GW144 |
| 1444 | NA | 27.2 | 1367.1 | NA | 2407.9 | 7849.6 | 477.4 | 472.9 | 285.6 | NA | 13.0 | 73.0 | 400.0 | 19484.0 | 7580.0 | 368.0 | 5.1 | 5.3 | 19.0 | 213.0 | 28.8 | NA | 216.0 | 2170.5 | 36.0 | 8.8 | 1177.5 | 119.0 | 9413.9 | 40.6 | 6.1 | 6.3 | 90 | 40 | GW144 |
| 1445 | NA | 19.8 | 1727.8 | NA | 3790.4 | 10221.7 | 713.3 | 1005.2 | 159.7 | NA | 12.0 | 84.0 | 71.0 | 9275.5 | 8463.0 | 245.0 | 6.2 | 4.5 | 18.0 | 125.0 | 21.8 | NA | 173.0 | 1230.0 | 31.0 | 7.6 | 875.5 | 94.0 | 13584.0 | 61.2 | 6.7 | 5.9 | 90 | 50 | GW144 |
| 1446 | NA | NA | 338.9 | NA | 507.7 | 791.5 | 108.7 | 44.2 | 172.1 | NA | NA | 63.0 | 67.0 | 20337.5 | 15651.0 | 553.0 | NA | 9.0 | 32.0 | 125.0 | 46.6 | NA | 153.0 | 2375.5 | 56.0 | 13.8 | 250.0 | 284.0 | 2642.8 | 10.5 | 1326.8 | 7.8 | 94 | 60 | GW144 |
| 1447 | 1.5 | NA | 652.4 | NA | 3329.0 | 753.3 | 118.3 | 354.4 | 226.9 | 25.0 | 12.0 | 121.0 | 403.0 | 50825.0 | 11032.0 | 675.0 | 8.2 | 6.8 | 106.0 | 209.0 | 34.6 | 82.0 | 181.0 | 1829.0 | 31.0 | 12.3 | 849.5 | 175.0 | 7350.1 | 33.0 | 4.2 | 7.5 | NA | 70 | GW144 |
| 531 | NA | NA | 560.3 | NA | 1547.8 | 1273.2 | 453.0 | 207.2 | 180.5 | 23.0 | 21.0 | 131.0 | 322.0 | 58495.0 | 10464.0 | 603.0 | NA | 4.1 | 43.0 | 140.0 | 30.0 | NA | 178.0 | 2113.0 | 29.0 | 8.2 | 575.5 | 191.0 | 4976.9 | 32.7 | 2.4 | 7.8 | 67.4 | 10 | GW53 |
| 532 | NA | NA | 586.0 | NA | 2678.0 | 1141.8 | 319.8 | 220.0 | 315.9 | 40.0 | 17.0 | 157.0 | 255.0 | 118809.5 | 8438.0 | 663.0 | 14.4 | NA | 86.0 | 146.0 | 24.6 | 226.0 | 219.0 | 2471.5 | NA | 6.7 | 665.0 | 121.0 | 5595.5 | 23.3 | 3.7 | 8.0 | 85.1 | 20 | GW53 |
| 533 | NA | NA | 672.3 | NA | 1393.5 | 1330.3 | 293.6 | 105.6 | 166.7 | NA | 19.0 | 55.0 | 81.0 | 31087.5 | 13030.0 | 1139.0 | NA | 9.9 | 29.0 | 173.0 | 58.1 | NA | 155.0 | 2282.5 | 47.0 | 12.3 | 295.5 | 242.0 | 2932.3 | 13.2 | 2.5 | 7.6 | 110 | 30 | GW53 |
| 534 | NA | NA | 436.2 | NA | 1811.3 | 1203.6 | 235.3 | 189.1 | 210.6 | NA | 30.0 | 77.0 | 113.0 | 21283.5 | 11394.0 | 1280.0 | NA | 9.6 | 34.0 | 205.0 | 37.5 | NA | 211.0 | 2346.5 | 42.0 | 12.6 | 928.5 | 233.0 | 3357.4 | 15.2 | 2.6 | 7.5 | 114.9 | 40 | GW53 |

| | | | | | | | | | | | | | | | | | | | | | | | | | | | | | | | | | | | |
|---|---|---|---|---|---|---|---|---|---|---|---|---|---|---|---|---|---|---|---|---|---|---|---|---|---|---|---|---|---|---|---|---|---|---|---|
| 535 | NA | NA | 1190.1 | NA | 2567.3 | 3853.7 | 530.2 | 226.9 | 394.2 | 20.0 | NA | 48.0 | 134.0 | 25309.0 | 11336.0 | 2235.0 | NA | 4.5 | 45.0 | 716.0 | 29.8 | 37.0 | 401.0 | 1803.5 | 70.0 | 7.9 | 1053.0 | 117.0 | 5992.7 | 29.7 | 4.6 | 6.8 | 111.3 | 50 | GW53 |
| 536 | NA | NA | 234.4 | NA | 2141.3 | 516.2 | 254.0 | 185.1 | 251.7 | NA | NA | 77.0 | 156.0 | 24674.0 | 12260.0 | 1523.0 | NA | 10.0 | 20.0 | 403.0 | 42.0 | NA | 260.0 | 2398.0 | 59.0 | 14.1 | 906.5 | 252.0 | 3317.5 | 14.8 | 2.4 | 7.8 | 114.2 | 60 | GW53 |
| 537 | NA | NA | 1601.1 | NA | 1524.9 | 3367.5 | 197.7 | 190.3 | 799.4 | NA | NA | 30.0 | 297.0 | 23635.5 | 7470.0 | 807.0 | NA | 4.3 | 31.0 | 138.0 | 18.5 | NA | 564.0 | 1558.0 | 19.0 | 4.0 | 507.0 | 102.0 | 4586.7 | 30.1 | 4.6 | 6.9 | 103.5 | 70 | GW53 |
| 538 | NA | NA | 302.4 | NA | 2840.3 | 588.9 | 169.2 | 255.1 | 257.1 | NA | 8.0 | 105.0 | 191.0 | 20118.0 | 8067.0 | 393.0 | NA | 4.9 | 44.0 | 94.0 | 22.9 | NA | 172.0 | 1847.5 | 25.0 | 7.3 | 609.5 | 138.0 | 5679.9 | 38.5 | 3.0 | 7.5 | 95 | 80 | GW53 |
| 539 | NA | 260.4 | 980.8 | NA | 2423.8 | 8942.6 | 754.7 | 330.3 | 187.8 | NA | NA | 76.0 | 1377.0 | 16653.0 | 5692.0 | 374.0 | NA | 3.1 | 50.0 | 671.0 | 18.2 | NA | 175.0 | 1575.5 | 32.0 | 5.5 | 2383.5 | 89.0 | 6065.0 | 33.6 | 6.2 | 6.5 | 93 | 88 | GW53 |
| 541 | 8.6 | 693.5 | 952.0 | 367.8 | 455.2 | 542.8 | 57.4 | 68.9 | 29.4 | NA | 21.0 | 98.0 | 219.0 | 48326.5 | 8393.0 | 404.0 | 11.0 | 5.2 | 41.0 | 136.0 | 23.7 | 56.0 | 313.0 | 2872.0 | 340.0 | 7.9 | 1097.5 | 144.0 | 5013.9 | 25.3 | 4.3 | 7.6 | 77 | 10 | GW54 |
| 542 | 6.6 | 689.5 | 284.9 | NA | 305.4 | 121.9 | 35.1 | 31.4 | 84.9 | NA | 19.0 | 60.0 | 110.0 | 25207.5 | 9707.0 | 424.0 | 4.5 | 7.2 | 20.0 | 171.0 | 31.9 | NA | 217.0 | 2233.0 | 42.0 | 9.9 | 447.5 | 221.0 | 5362.4 | 18.1 | 1.9 | 8.3 | 85 | 20 | GW54 |
| 543 | 6.5 | NA | 737.5 | NA | 397.9 | 382.6 | 44.1 | 59.7 | 135.9 | NA | 13.0 | 47.0 | 2534.0 | 31094.5 | 12600.0 | 359.0 | NA | 5.0 | 20.0 | 129.0 | 32.4 | 39.0 | 253.0 | 1811.5 | 23.0 | 10.8 | 1879.0 | 157.0 | 4672.8 | 19.5 | 3.5 | 7.8 | 96 | 30 | GW54 |
| 544 | 6.3 | NA | 108.9 | 365.6 | 247.6 | 69.9 | 12.6 | 28.8 | 102.6 | 27.0 | NA | 88.0 | 149.0 | 35673.0 | 12968.0 | 735.0 | NA | 9.1 | 35.0 | 208.0 | 43.7 | 32.0 | 186.0 | 2737.0 | 43.0 | 13.4 | 1840.0 | 286.0 | 3306.5 | 13.7 | 1.7 | 7.9 | 118 | 40 | GW54 |
| 545 | 4.8 | 694.4 | 626.7 | 365.1 | 579.2 | 373.7 | 64.2 | 95.5 | 188.1 | NA | NA | 786.0 | 374.0 | 29214.5 | 8005.0 | 482.0 | NA | NA | NA | 106.0 | 18.5 | NA | 143.0 | 2181.5 | 102.0 | 5.9 | 19494.5 | 98.0 | 7630.9 | 40.5 | 4.6 | 6.7 | 104 | 50 | GW54 |
| 546 | 6.5 | 691.5 | 999.5 | NA | 228.2 | 706.2 | 43.3 | 33.7 | 43.0 | NA | 14.0 | 47.0 | 253.0 | 21730.0 | 7198.0 | 523.0 | NA | 4.4 | 27.0 | 98.0 | 23.9 | NA | 412.0 | 1853.5 | 31.0 | 9.1 | 950.5 | 170.0 | 4022.1 | 22.0 | 3.4 | 7.4 | 103 | 60 | GW54 |
| 547 | 5.6 | NA | 162.4 | NA | 424.4 | 118.4 | 16.0 | 38.5 | 77.5 | 26.0 | 20.0 | 54.0 | 184.0 | 35895.5 | 11048.0 | 533.0 | NA | 7.2 | 34.0 | 201.0 | 36.4 | 51.0 | 278.0 | 2757.0 | 39.0 | 9.8 | 854.0 | 194.0 | 4306.2 | 20.1 | 2.5 | 8.5 | 122 | 70 | GW54 |
| 548 | 6.5 | 695.4 | 1359.0 | NA | 184.1 | 849.7 | 59.5 | 31.6 | 58.4 | NA | NA | 48.0 | 317.0 | 22340.0 | 11839.0 | 459.0 | NA | 9.4 | 26.0 | 103.0 | 37.7 | NA | 248.0 | 2570.0 | 37.0 | 11.2 | 982.0 | 223.0 | 3970.8 | 13.4 | 3.0 | 8.0 | 103 | 80 | GW54 |
| S10 | 0.4 | NA | 98.0 | NA | 46.3 | 360.7 | 48.2 | 48.2 | 5.0 | NA | 8.9 | 42.0 | 22.0 | 17955.5 | 16224.0 | 582.0 | NA | 13.2 | 26.0 | 23.0 | 60.9 | NA | 127.0 | 2694.5 | 61.0 | 20.4 | 61.5 | 385.0 | 2311.0 | 3.2 | 0.3 | 8.1 | NA | 3 | Surface |
| S6 | 0.4 | NA | 234.0 | NA | 69.9 | 705.4 | 69.9 | 69.9 | 9.1 | NA | 7.3 | 38.0 | 20.0 | 15845.0 | 15129.0 | 562.0 | NA | 12.5 | 26.0 | 22.0 | 54.4 | NA | 121.0 | 2234.0 | 54.0 | 18.3 | 46.5 | 352.0 | 2934.0 | 2.3 | 0.1 | 7.9 | NA | 3 | Surface |
| S7 | 0.3 | NA | 168.7 | NA | 89.4 | 465.8 | 59.8 | 59.8 | 4.2 | NA | 6.7 | 43.0 | 19.0 | 18710.0 | 15672.0 | 607.0 | NA | 12.4 | 27.0 | 18.0 | 55.4 | NA | 122.0 | 2389.5 | 54.0 | 20.7 | 45.5 | 344.0 | 1174.0 | 1.9 | 0.4 | 8.0 | NA | 3 | Surface |
| S8 | 0.6 | NA | 140.9 | NA | 217.9 | 508.4 | 68.2 | 68.2 | 8.3 | NA | 9.1 | 43.0 | 22.0 | 18633.0 | 14962.0 | 557.0 | NA | 12.2 | 30.0 | 34.0 | 61.8 | NA | 126.0 | 2737.5 | 67.0 | 20.9 | 68.5 | 387.0 | 1616.3 | 3.3 | 0.6 | 8.0 | NA | 3 | Surface |
| S9 | 1.5 | NA | 154.5 | NA | 125.9 | 506.9 | 106.9 | 106.9 | 10.5 | NA | 7.8 | 29.0 | 18.0 | 16579.0 | 15092.0 | 540.0 | NA | 9.8 | 21.0 | 18.0 | 53.2 | NA | 120.0 | 2424.5 | 49.0 | 17.7 | 49.0 | 327.0 | 1273.2 | 3.0 | 0.5 | 8.0 | NA | 3 | Surface |
| SC1 | 1.1 | NA | 7.4 | 0.1 | 24.0 | 120.5 | 21.9 | 38.2 | 2.5 | 25.0 | 6.9 | 31.7 | 21.3 | 13833.8 | 16604.3 | 594.0 | NA | 9.3 | 25.7 | 13.8 | 48.3 | NA | 118.7 | 2001.0 | 42.3 | 15.9 | 35.2 | 329.7 | 1671.2 | 1.7 | 0.1 | 8.7 | NA | Control | Control |

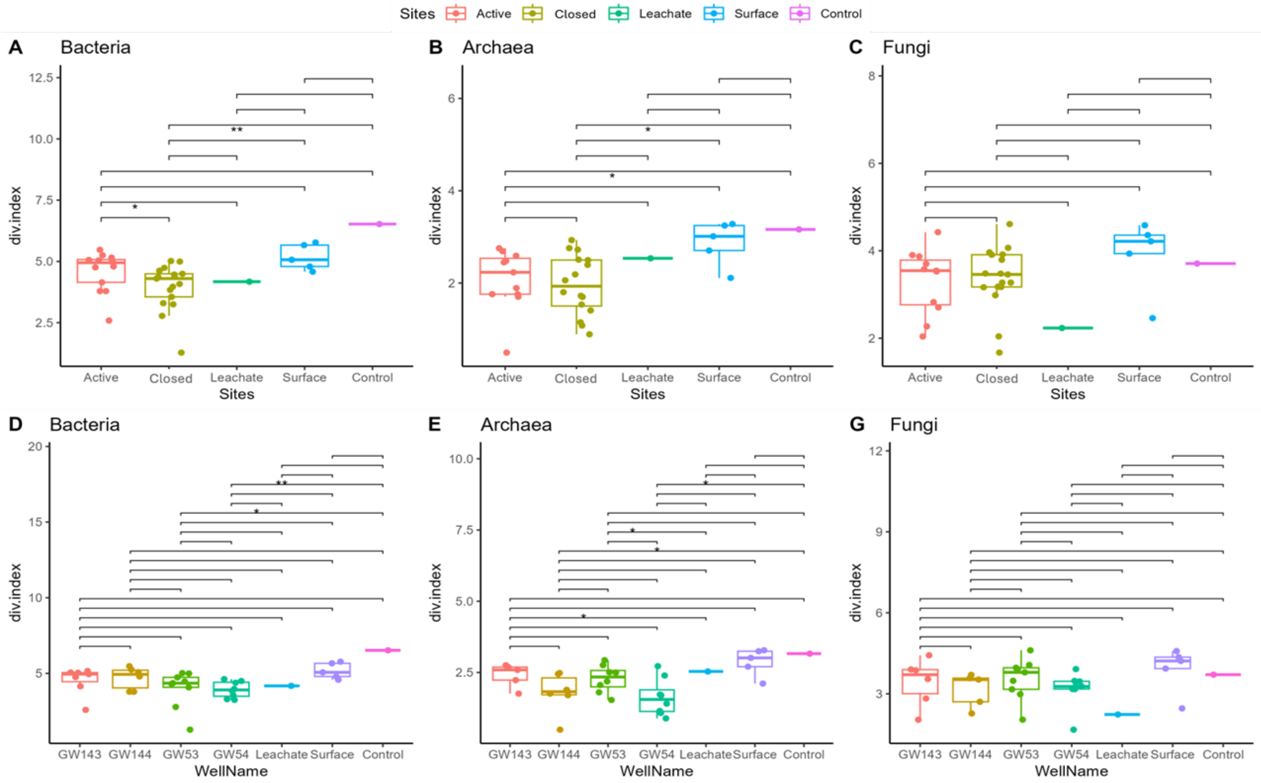


Figure S3. Alpha diversity of bacterial, archaeal, and fungal communities across landfill sites and wells. Boxplots show diversity indices for bacteria (A, D), archaea (B, E), and fungi (C, G) based on observed richness (A–C) and Shannon diversity index (D–G) across five site types: Active, Closed, Leachate, Surface, and Control. Panels A–C display site-level comparisons, while panels D–G show well-level variation, including wells GW143, GW144 (Active), GW53, GW54 (Closed), and other site types. Distinct colors represent different site categories. Asterisks (*) indicate significant pairwise differences ($p < 0.05$); “ns” indicates non-significant comparisons.

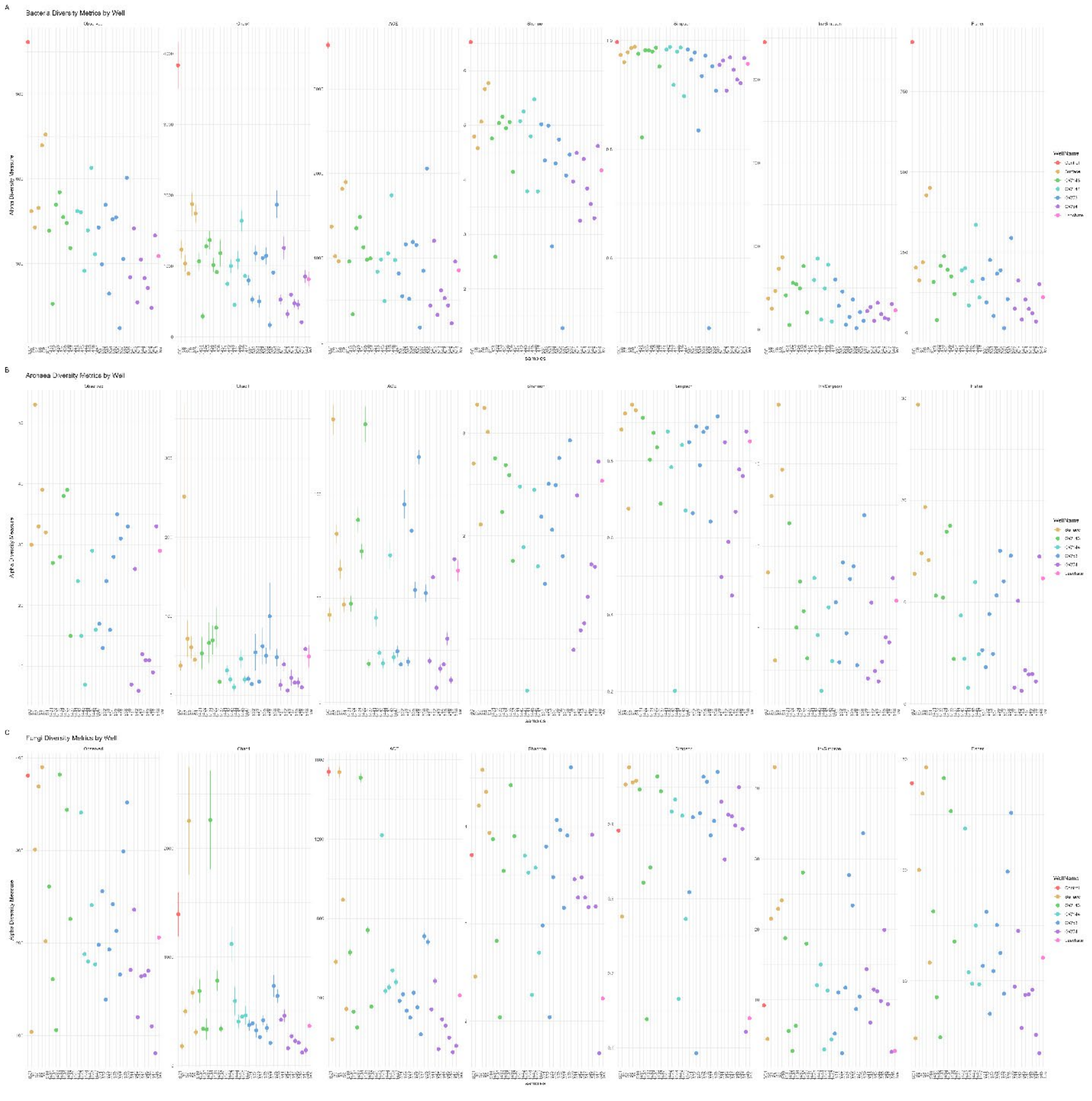


Figure S4. Alpha diversity metrics for microbial communities (A)bacteria, (B) archaea, and (C) fungi) across different landfill samples. The metrics displayed include Chao1 (species richness estimator), Observed (number of OTU's - unique taxa), ACE (abundance-based coverage estimator), Shannon (diversity index accounting for richness and evenness), Simpson (dominance measure of common species), and Fisher's Alpha (diversity estimator based on species abundance distribution). Each point represents a sample, color-coded by category. The x-axis indicates different sample groups, while the y-axis displays the diversity values for each metric.

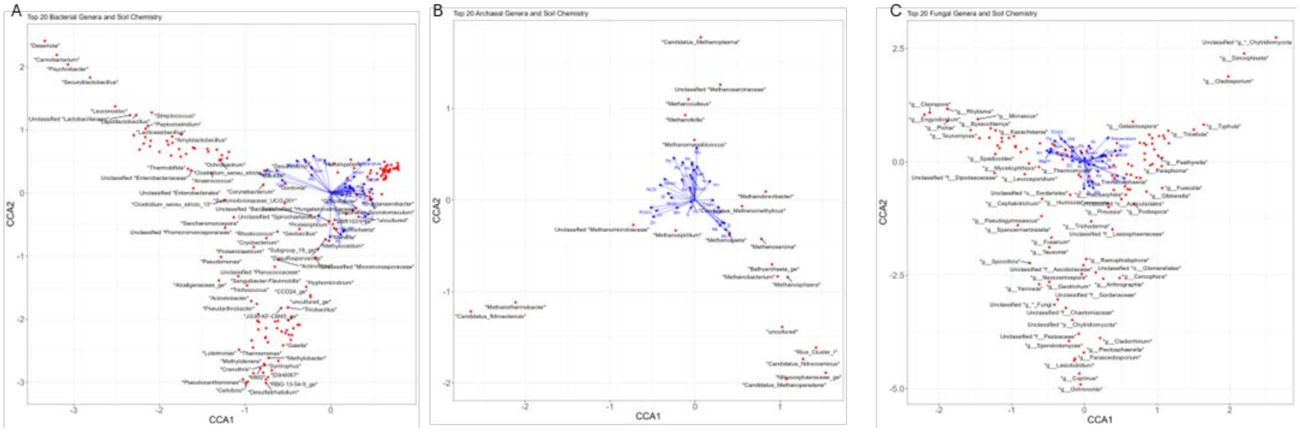


Figure S5. Canonical Correspondence Analysis (CCA) of microbial genera and soil chemistry across landfill samples. (A) Bacterial genera, (B) Archaeal genera, and (C) Fungal genera. Each plot displays the top 20 most abundant genera and their ordination relative to environmental variables (blue arrows), including pH, sulfate, ammonia, nitrate, and others. Red points represent individual genera. The distinct clustering and orientation of taxa and samples suggest that bacteria, archaea, and fungi are each shaped by different combinations of environmental factors, highlighting niche specialization across microbial domains.

Table S5: Methane, Oxygen and Temperature composition over time in landfills wells

| Point Name | Sample Date | Methane (%) | Oxygen (%) | Temperature (F) |
|---|---|---|---|---|
| GW-53 | 5/24/2021 | 45.7 | 1.9 | 104.7 |
| GW-54 | 5/24/2021 | 54.6 | 0 | 86.2 |
| GW-143 | 6/29/2021 | 38.1 | 0 | 105.3 |
| GW-144 | 6/29/2021 | 40.9 | 0 | 113.4 |
| GW-53 | 6/30/2021 | 47.5 | 1.7 | 107.2 |
| GW-54 | 6/30/2021 | 53.8 | 0 | 113.2 |
| GW-53 | 7/29/2021 | 41.7 | 3.7 | 100.6 |
| GW-54 | 7/29/2021 | 50 | 1.8 | 113.7 |
| GW-143 | 7/29/2021 | 46.4 | 0 | 104.5 |
| GW-144 | 7/29/2021 | 51.1 | 0 | 113.4 |
| GW-143 | 8/24/2021 | 48.6 | 0 | 107.1 |
| GW-144 | 8/24/2021 | 55.2 | 0 | 113.5 |
| GW-54 | 8/24/2021 | 55.3 | 0 | 110.3 |
| GW-53 | 8/24/2021 | 56.9 | 0 | 120.4 |
| GW-54 | 9/14/2021 | 54.2 | 0.3 | 108.5 |
| GW-53 | 9/14/2021 | 57.4 | 0 | 115.3 |
| GW-143 | 9/15/2021 | 47.6 | 0 | 107.1 |
| GW-144 | 9/15/2021 | 49.9 | 0 | 114.4 |
| GW-54 | 10/18/2021 | 46.5 | 2.7 | 72.9 |
| GW-144 | 10/18/2021 | 51.5 | 0 | 112.3 |
| GW-143 | 10/18/2021 | 51.7 | 0 | 106.7 |
| GW-53 | 10/18/2021 | 59.1 | 0 | 108 |
| GW-144 | 11/23/2021 | 51.8 | 0 | 112.6 |
| GW-143 | 11/23/2021 | 52.8 | 0 | 105.6 |
| GW-54 | 11/23/2021 | 56.8 | 0 | 98.8 |
| GW-53 | 11/23/2021 | 58.7 | 0 | 106 |
| GW-143 | 12/16/2021 | 50.1 | 0 | 104.9 |
| GW-144 | 12/16/2021 | 52.5 | 0 | 111.9 |
| GW-54 | 12/16/2021 | 55.6 | 0 | 47.3 |
| GW-53 | 12/16/2021 | 56.8 | 0 | 102.7 |
| GW-53 | 1/17/2022 | 58.5 | 0 | 102.4 |
| GW-144 | 1/18/2022 | 51.2 | 0 | 112.6 |
| GW-143 | 1/18/2022 | 51.7 | 0 | 105.8 |
| GW-54 | 1/18/2022 | 53.6 | 0 | 104.7 |
| GW-54 | 2/10/2022 | 47.3 | 3.3 | 95.2 |
| GW-144 | 2/10/2022 | 51.7 | 0 | 113 |
| GW-143 | 2/10/2022 | 53.4 | 0 | 106.7 |
| GW-53 | 2/10/2022 | 59.4 | 0 | 102.4 |
| GW-54 | 3/1/2022 | 49.3 | 1.4 | 108.7 |

| | | | | |
|---|---|---|---|---|
| GW-144 | 3/1/2022 | 52.2 | 0 | 113.2 |
| GW-143 | 3/1/2022 | 53.3 | 0 | 108.9 |
| GW-53 | 3/1/2022 | 59.4 | 0 | 99.9 |
| GW-144 | 4/26/2022 | 50.4 | 0 | 115 |
| GW-143 | 4/26/2022 | 52.3 | 0 | 108.5 |
| GW-54 | 4/26/2022 | 53.7 | 0 | 118.4 |
| GW-53 | 4/26/2022 | 54 | 0 | 103.1 |
| GW-144 | 5/19/2022 | 46.4 | 0 | 114.8 |
| GW-143 | 5/19/2022 | 50.6 | 0 | 110.3 |
| GW-53 | 5/19/2022 | 53 | 0 | 110.7 |
| GW-54 | 5/19/2022 | 53.7 | 0 | 116.8 |
| GW-144 | 6/23/2022 | 47.3 | 0 | 115.5 |
| GW-143 | 6/23/2022 | 50.3 | 0 | 112.6 |
| GW-54 | 6/23/2022 | 50.4 | 0 | 114.4 |
| GW-53 | 6/23/2022 | 51.8 | 0 | 109.8 |
| GW-144 | 7/20/2022 | 47.1 | 0 | 115 |
| GW-53 | 7/20/2022 | 50.6 | 0 | 109.9 |
| GW-143 | 7/20/2022 | 50.9 | 0 | 113 |
| GW-54 | 7/20/2022 | 52.2 | 0 | 109.6 |
| GW-144 | 8/4/2022 | 46.4 | 0 | 117 |
| GW-54 | 8/4/2022 | 51.7 | 0 | 113 |
| GW-143 | 8/4/2022 | 52 | 0 | 113.7 |
| GW-53 | 8/4/2022 | 53.2 | 0 | 109.8 |
| GW-144 | 9/15/2022 | 47.9 | 0 | 116.6 |
| GW-143 | 9/15/2022 | 52.8 | 0 | 113.4 |
| GW-53 | 9/15/2022 | 53.4 | 0 | 109.8 |
| GW-54 | 9/15/2022 | 56.7 | 0 | 110.3 |
| GW-144 | 10/27/2022 | 47.1 | 0 | 104.7 |
| GW-53 | 10/27/2022 | 53 | 0 | 108.3 |
| GW-143 | 10/27/2022 | 53.9 | 0 | 115.9 |
| GW-54 | 10/27/2022 | 54.8 | 0 | 108.5 |
| GW-143 | 11/28/2022 | 47.9 | 0 | 114.1 |
| GW-144 | 11/28/2022 | 50.5 | 0 | 104.5 |
| GW-53 | 11/28/2022 | 51.5 | 0 | 104.5 |
| GW-54 | 11/28/2022 | 52.4 | 0 | 104.9 |
| GW-144 | 12/12/2022 | 46.4 | 0 | 92.3 |
| GW-143 | 12/12/2022 | 47.5 | 0 | 113.9 |
| GW-53 | 12/12/2022 | 51.4 | 0 | 107.1 |
| GW-54 | 12/12/2022 | 53 | 0 | 106.9 |
| GW-143 | 1/17/2023 | 48.4 | 0 | 114 |
| GW-144 | 1/17/2023 | 48.4 | 0 | 114 |

| | | | | |
|---|---|---|---|---|
| GW-53 | 1/17/2023 | 56 | 0 | 100 |
| GW-54 | 1/17/2023 | 56.6 | 0 | 100 |
| GW-143 | 2/13/2023 | 47.8 | 0 | 112 |
| GW-144 | 2/13/2023 | 48.7 | 0 | 114 |
| GW-53 | 2/13/2023 | 53.8 | 0 | 102 |
| GW-54 | 2/13/2023 | 55.5 | 0 | 102 |
| GW-144 | 3/13/2023 | 47.7 | 0 | 85.6 |
| GW-143 | 3/13/2023 | 49.4 | 0 | 88.7 |
| GW-53 | 3/13/2023 | 52.6 | 0 | 99.1 |
| GW-54 | 3/13/2023 | 55.1 | 0 | 102.2 |
| GW-143 | 4/17/2023 | 47.2 | 0 | 106 |
| GW-144 | 4/17/2023 | 47.3 | 0 | 86.2 |
| GW-53 | 4/17/2023 | 52.4 | 0 | 100.4 |
| GW-54 | 4/17/2023 | 54.6 | 0 | 103.3 |
| GW-143 | 5/15/2023 | 48.3 | 0 | 104.2 |
| GW-144 | 5/15/2023 | 49.3 | 0 | 108.5 |
| GW-54 | 5/15/2023 | 52.9 | 0 | 104.5 |
| GW-53 | 5/15/2023 | 54.4 | 0 | 102.2 |
| GW-54 | 6/12/2023 | 43 | 2.9 | 106.3 |
| GW-144 | 6/12/2023 | 44.4 | 0 | 112.8 |
| GW-143 | 6/12/2023 | 48 | 0 | 119.7 |
| GW-53 | 6/12/2023 | 54.1 | 0 | 107.6 |
| GW-143 | 7/18/2023 | 46.9 | 0 | 119.5 |
| GW-144 | 7/18/2023 | 47.9 | 0 | 80.6 |
| GW-53 | 7/18/2023 | 52.9 | 0 | 107.8 |
| GW-54 | 7/18/2023 | 54.6 | 0 | 106.3 |
| GW-143 | 8/17/2023 | 48.2 | 0 | 120.9 |
| GW-144 | 8/17/2023 | 48.5 | 0 | 114.8 |
| GW-53 | 8/17/2023 | 53.5 | 0 | 106.3 |
| GW-54 | 8/17/2023 | 54.6 | 0 | 104.9 |
| GW-144 | 9/12/2023 | 45.1 | 0 | 115.3 |
| GW-143 | 9/12/2023 | 48.3 | 0 | 120.2 |
| GW-54 | 9/12/2023 | 55.4 | 0 | 105.6 |
| GW-53 | 9/12/2023 | 55.9 | 0 | 106.5 |
| GW-143 | 10/16/2023 | 47.2 | 0 | 120 |
| GW-144 | 10/16/2023 | 48.9 | 0 | 90.9 |
| GW-53 | 10/16/2023 | 54.3 | 0 | 100.9 |
| GW-54 | 10/16/2023 | 55.1 | 0 | 104.7 |
| GW-143 | 11/13/2023 | 42.1 | 0.6 | 117.1 |
| GW-144 | 11/13/2023 | 49.3 | 0 | 118.9 |
| GW-54 | 11/13/2023 | 54.3 | 0 | 97.7 |

| | | | | |
|---|---|---|---|---|
| GW-53 | 11/13/2023 | 54.8 | 0 | 101.5 |
| GW-143 | 12/11/2023 | 44.8 | 0 | 119.7 |
| GW-144 | 12/11/2023 | 51.1 | 0 | 116.8 |
| GW-53 | 12/11/2023 | 55.3 | 0 | 100.6 |
| GW-54 | 12/11/2023 | 55.5 | 0 | 102.7 |
| GW-53 | 1/10/2024 | 54.5 | 0 | 100.9 |
| GW-54 | 1/10/2024 | 55.3 | 0 | 99.3 |
| GW-144 | 1/11/2024 | 47.3 | 0 | 115.2 |
| GW-143 | 1/11/2024 | 49.2 | 0 | 113.2 |
| GW-143 | 2/19/2024 | 51.4 | 0 | 110.1 |
| GW-144 | 2/19/2024 | 51.4 | 0 | 118.6 |
| GW-53 | 2/19/2024 | 53.5 | 0 | 101.7 |
| GW-54 | 2/19/2024 | 55.4 | 0 | 99 |
| GW-144 | 3/18/2024 | 50.7 | 0 | 117.3 |
| GW-143 | 3/18/2024 | 52 | 0 | 118.9 |
| GW-53 | 3/18/2024 | 55 | 0 | 98.6 |
| GW-54 | 3/18/2024 | 57.5 | 0 | 92.1 |
| GW-143 | 4/15/2024 | 49 | 0.1 | 123.8 |
| GW-144 | 4/15/2024 | 49.9 | 0.1 | 117.7 |
| GW-53 | 4/15/2024 | 54.4 | 0.1 | 98.8 |
| GW-54 | 4/15/2024 | 56.4 | 0.1 | 93 |
| GW-53 | 5/20/2024 | 56.7 | 0.1 | 97.2 |
| GW-54 | 5/20/2024 | 56.9 | 0.1 | 85.1 |
| GW-143 | 5/21/2024 | 50.6 | 0 | 122 |
| GW-144 | 5/21/2024 | 50.6 | 0 | 120.7 |
| GW-53 | 6/24/2024 | 53.3 | 0.6 | 100 |
| GW-144 | 6/24/2024 | 45.3 | 0.1 | 121.3 |
| GW-143 | 6/24/2024 | 47.8 | 0.1 | 124.7 |
| GW-54 | 6/24/2024 | 56.8 | 0.1 | 90.5 |
| GW-144 | 7/22/2024 | 46.7 | 0.1 | 123.4 |
| GW-143 | 7/22/2024 | 50.7 | 0.1 | 123.8 |
| GW-54 | 7/22/2024 | 56.6 | 0.1 | 94.8 |
| GW-53 | 7/22/2024 | 57 | 0.1 | 101.8 |
| GW-54 | 8/19/2024 | 55.7 | 0.1 | 94.1 |
| GW-143 | 8/19/2024 | 52.1 | 0 | 123.1 |
| GW-144 | 8/19/2024 | 54.6 | 0 | 122.7 |
| GW-53 | 8/19/2024 | 56.9 | 0 | 101.1 |
| GW-144 | 9/12/2024 | 43.3 | 0.1 | 122.4 |
| GW-54 | 9/12/2024 | 55.1 | 0.1 | 96.1 |
| GW-53 | 9/12/2024 | 55.4 | 0.1 | 104 |
| GW-143 | 9/13/2024 | 52.7 | 0.2 | 123.4 |

Landfill environmental monitoring data, such as release of oxygen, methane, hydrogen, among others, were obtained from Waste & Materials Management GEMS on the Web (GOTW) Public Access (https://apps.dnr.wi.gov/gotw/webpages/UserAgreement.aspx)
